# A reversible YbbR-domain hinge switch senses extracellular pH to control bacterial cyclic-di-AMP signaling

**DOI:** 10.64898/2026.11.11.765923

**Authors:** Katharina Weinhäupl, Rajkumar Singh, Teresa Megre Lousada, Arantza Muguruza-Montero, Isabel M. Matas, Hugo Fraga, Paul Schanda, João H. Morais-Cabral

## Abstract

Acidification is a principal antimicrobial host defence: during infection, bacteria have to survive the stomach, the vaginal tract or the phagolysosome of immune cells, where the pH falls below 5. To persist, bacteria must sense and adapt to extracellular pH. In Gram-positive bacteria, the membrane protein CdaR, a regulator of the c-di-AMP-synthesizing enzyme CdaA, has been proposed as a pH sensor, but the molecular basis of this sensing remains unknown.

Here we show, using the model organism *Bacillus subtilis*, that CdaR is essential for growth below pH 5. SEC-MALS, mass photometry and circular dichroism demonstrate that its extracellular YbbR domains neither oligomerize nor unfold at low pH. Instead, tryptophan and ANS fluorescence reveal a reversible conformational change that requires at least two tandem YbbR domains and occurs between pH 4.5 and 5.5. Crystallography, molecular dynamics and solution NMR localize this transition to a conserved hinge between YbbR domains 1 and 2, where a network of charged residues rearranges to rigidify it and restrict inter-domain motion. Mutagenesis identifies a highly conserved aspartate whose substitution abolishes the conformational switch *in vitro* and accelerates adaptation to low pH *in vivo*.

These findings define a structural mechanism by which CdaR senses extracellular acidification through its YbbR domains and couples this signal to intracellular c-di-AMP signaling. As the hinge and its aspartate are the most conserved features of the YbbR family, found in pathogens including *Staphylococcus aureus* and *Listeria monocytogenes*, we propose this as a general mechanism of acid adaptation in Gram-positive bacteria.

## Introduction

As unicellular organisms, bacteria need to adapt rapidly and efficiently to changes in their environment. To this end, prokaryotes, as well as eukaryotes, use second messenger systems to regulate a multitude of cellular processes. The second messenger cyclic-di-AMP (c-di-AMP), discovered only in 2008(1), is exclusively found in prokaryotes and has since been implicated in DNA integrity surveillance, cell wall homeostasis, and sporulation. It is essential for the survival and virulence of many Gram-positive bacteria. Intracellular levels of this second messenger are tightly regulated by proteins that either synthesize or degrade c-di-AMP. All Gram-positive bacteria express at least one diadenylate cyclase for synthesis and one phosphodiesterase for degradation(2).

One environmental challenge frequently faced by bacteria is fluctuations in pH(3–5). While the majority of species exhibit optimal growth at or near pH neutrality, bacteria will encounter more extreme environments with pH values below 5. In natural ecosystems, acidic soils or peatlands can reach pH values between 3 - 5. Similarly, food products such as fruits, fruit juices, and fermented products typically fall within the range of pH 3 - 4. Within an animal host, bacteria must withstand strong acidity in the gastric compartment (pH 1 - 3, rising to 4 - 5 postprandially), the vaginal tract (pH 3.5 - 4.5), and the phagosomal/lysosomal pathway of immune cells, where the pH decreases to approximately 4.5 - 5 as part of the antimicrobial response (reviewed in (6)).

To survive under such conditions, bacteria have developed multiple acid-resistance mechanisms: Protons can be actively extruded via the F₀F₁-ATPase and secondary antiporters such as K⁺/H⁺ and Na⁺/H⁺ exchangers(7). Intracellular pH homeostasis is further maintained by metabolic buffering systems. *Helicobacter pylori* deploys a potent urease that generates ammonia from urea(8), whereas enterobacteria activate amino acid decarboxylase systems (glutamate, arginine, lysine), in which decarboxylation consumes cytosolic protons and the resulting products are exchanged for new substrate via cognate antiporters(9). Acid shock proteins, including the periplasmic chaperones HdeA and HdeB, prevent acid-induced protein aggregation(10), while DNA-binding proteins such as Dps and HU protect genomic integrity(11, 12). Global stress regulators, notably σ^S^ (RpoS) in Gram-negative bacteria and σ^B^ in Gram-positive species, coordinate the induction of a broad array of acid stress response genes(13, 14).

In host-associated environments, pathogens have evolved additional strategies to withstand acidity. *Mycobacterium tuberculosis* and *Salmonella enterica* can modulate phagosomal acidification(15, 16), while *Listeria monocytogenes* uses the acidic phagosomal environment to escape into the host cytosol before lysosomal fusion(17).

Biofilm formation provides extracellular buffering and creates localized microenvironments that are less susceptible to acidification(18, 19). Spore-forming taxa such as *Bacillus* and *Clostridium* survive extreme acidity in dormant spore states until favorable conditions are restored(20, 21). Collectively, these physiological, biochemical, and structural adaptations enable bacteria not only to survive but, in some cases, to proliferate in environments with pH well below 5, illustrating the remarkable plasticity of microbial stress responses.

To be able to react to extracellular acid stress, bacteria have evolved dedicated pH-sensing systems(22). In Gram-negative bacteria, several systems are known, including PhoQ/PhoP(23) and the antiporter GadC(24). In Gram-positive bacteria, a possible candidate for extracellular pH-sensing is the CdaR protein of the CdaA-CdaR-GlmM complex.

CdaA is a membrane-associated protein containing an intracellular diadenylate cyclase (DAC) domain responsible for c-di-AMP synthesis(25). CdaA has been implicated in various cellular processes, including potassium homeostasis(26), biofilm formation(27), and antibiotic resistance(28). In *Bacillus subtilis*, the cda-glm operon additionally encodes two regulatory proteins: the membrane protein CdaR and the cytosolic protein GlmM (Figure 1A). GlmM is involved in cell wall synthesis and interacts with the intracellular catalytic domain of CdaA, inhibiting its activity(29). Genetic and biochemical studies in various species have demonstrated that CdaR modulates the activity of CdaA: in some contexts, it inhibits, whereas in others it stimulates c-di-AMP synthesis, suggesting a context-dependent regulatory mechanism(25, 30).

**Figure 1.**
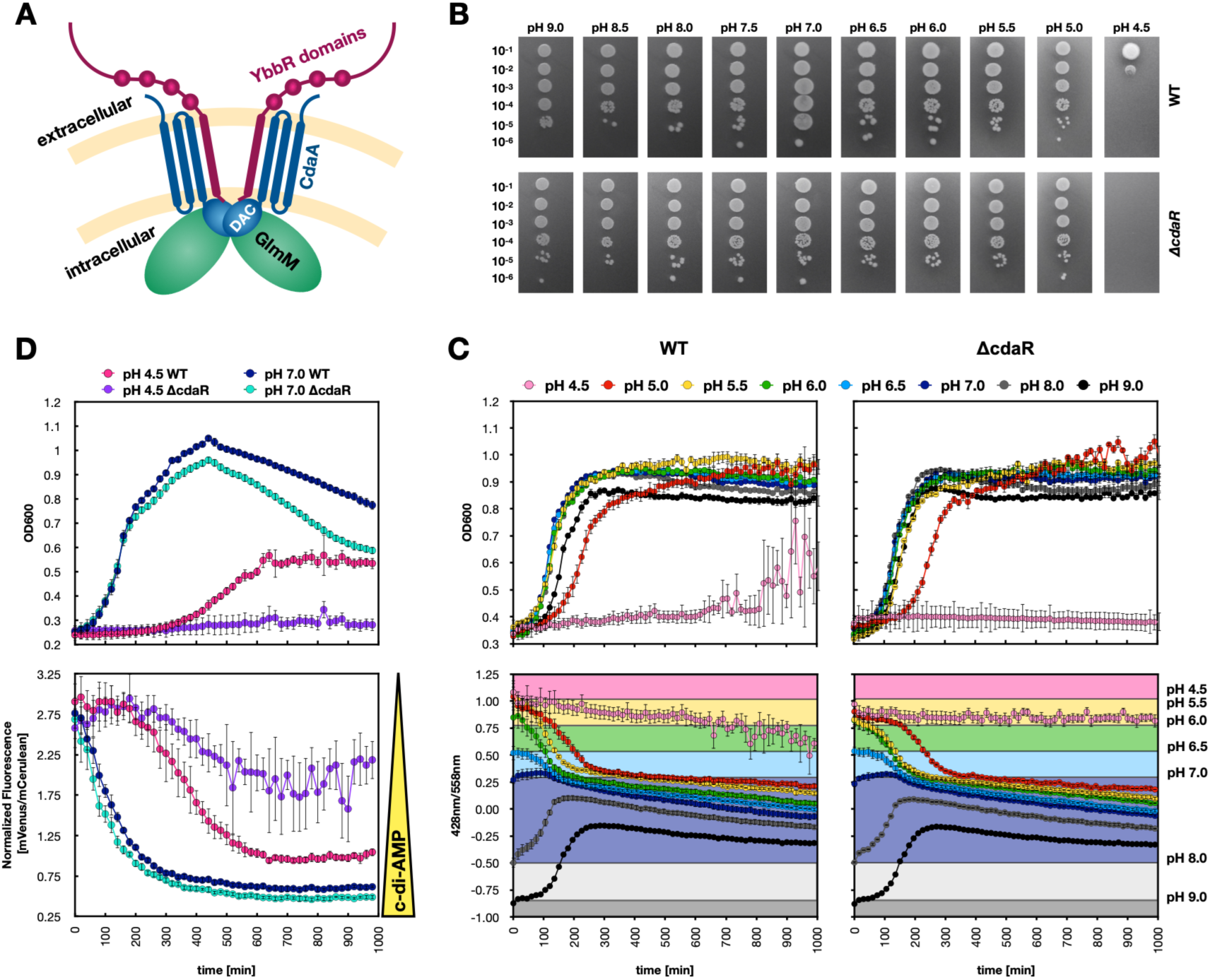
CdaR is essential for *Bacillus subtilis* growth at low pH. A) Graphical description of the CdaA-CdaR-GlmM complex. The activity of the diadenylate cyclase CdaA (in blue), is influenced by two regulatory proteins, the cytosolic GlmM (in green) and the membrane protein CdaR (in purple). B) Spot assay with dilution series of *B. subtilis* WT and Δ*cdaR* mutant on LB_BS_ Agar at different pHs. C) Growth of *B. subtilis* WT (left) and Δ*cdaR* (right) in LB_BS_ media with pH values ranging from pH 4.5 - 9.0 with simultaneous measurement of the external pH using the colorimetric pH sensor phenol red (bottom). D) Top: Growth curves of *B. subtilis* WT and Δ*cdaR* in minimal medium at pH 7.0 and 4.5. Bottom: cyclic-di-AMP production of *B. subtilis* WT and ΔcdaR at pH 7.0 and 4.5.

In *Staphylococcus aureus*, c-di-AMP levels increase at lower pH when CdaR is removed, suggesting a role in pH sensing(31). Somehow the cell must sense extracellular pH and transmit this information across the membrane to control intracellular c-di-AMP synthesis by CdaA. An obvious candidate for this sensing mechanism is the extracellular array of YbbR domains present in CdaR.

CdaR is a predominantly extracellular protein composed of a series of globular YbbR domains arranged like beads on a string and encoded in the same operon as CdaA. CdaR typically comprises a transmembrane helix and two to four tandem YbbR domains exposed to the periplasm or cell wall environment. In some bacterial species, hybrid proteins exist that combine the features of both CdaA and CdaR within a single polypeptide. YbbR domains constitute a conserved bacterial protein family distributed across diverse phyla. Although their primary sequence identity is low, they adopt a structurally conserved β-rich fold(32). Crystal structures of isolated YbbR domains from *Streptococcus pneumoniae* (PDB: 4QDY) and *B. subtilis* (PDB: 2O3F) have revealed a distinctive topology characterized by β-strands looping around the domain core, a conserved cis-proline residue stabilized by conserved valine residues.

The number of YbbR domains, as well as the presence or absence of intrinsically disordered regions (IDRs), varies substantially between species. In some organisms, CdaR contains IDRs appended to the YbbR repeats that are required for viability during envelope stress(33). These regions exhibit low sequence conservation but share compositional similarities with other IDRs known to interact with the *B. subtilis* cell wall(34). They are thought to penetrate gaps in the peptidoglycan meshwork or interact non-covalently with the cell wall and, together with the YbbR domains, transmit envelope perturbations to the cell interior. However, the mechanism by which signals from the cell wall are relayed through CdaR to CdaA remains unknown. Importantly, many CdaR homologues do not contain IDRs, among them well-studied organisms in the context of c-di-AMP signaling, such as *S. aureus* and *Lactobacillus acidophilus*. This observation raises the possibility that CdaR possesses functions beyond envelope damage sensing.

Here, we investigate the function of the YbbR domains of CdaR using the model organism *B. subtilis* and aim to uncover the molecular mechanism underlying extracellular pH sensing. At present, there is no structural or functional understanding of how YbbR domains sense environmental changes and transmit signals across the membrane. Do the YbbR domains undergo a conformational switch, oligomerize, or recruit additional factors that regulate c-di-AMP production?

## Results

### CdaR is essential for *Bacillus subtilis* growth at low pH

A growth defect of a ΔcdaR mutant at low pH has been previously described for *S. aureus*(*31*) and we aimed to confirm this phenotype in the *B. subtilis* strain 168. As before, deleting *cdaR* in *B. subtilis* leads to a growth defect in liquid and solid medium below pH 5 (Figure 1B/C). While growth at pH 5 is similar in the WT and Δ*cdaR* mutant, at pH 4.5 no growth of the mutant occurs within 24h at 37°C. We noted several phenotypic differences between the wild type and mutant strains. Using the pH sensor phenol red we found that WT *B. subtilis* is able to modify its environment during growth so that after the exponential phase all cultures had reached a pH in the range of 7.0 - 8.0, when the starting pH of the media was between 5 and 9 (Figure 1C). The wild type culture started at pH 4.5 shows a tendency for a change towards higher pH but did not reach the plateau phase at the end of the experiment. In contrast, in pH 4.5, the Δ*cdaR* mutant did not show any change in the external pH of the growth medium (Figure 1C, right), indicating that the environmental modification is linked to cell viability and growth. Additionally, images taken of *B. subtilis* WT and Δ*cdaR* after 16h growth at pH 7 or pH 4.5 show distinct phenotypes (Fig. S1A). At pH 7 the mutant and the WT do not show appreciable differences. In contrast, at pH 4.5 the WT, which has grown to the same cell density as at pH 7, grows in long filaments while the Δ*cdaR* mutant, which has shown no cell growth during this period, forms smaller granulated cells. We wondered if these structures are endospores and performed Malachite green staining to confirm this hypothesis; however, none of the bright structures showed a positive endospore stain (Fig. S1B).

To explore if the effects caused by deletion of *cdaR* are correlated with changes in internal c-di-AMP level, we investigated the impact of low pH on the production of c-di-AMP in *B. subtilis*. To this end we optimized a previously described c-di-AMP biosensor based on the *kimA* riboswitch(35). In the absence of c-di-AMP a fluorescent protein (mVenus) is transcribed and the fluorescence at 508 nm is high; when c-di-AMP levels increase, it binds to the riboswitch and terminates the transcription of the fluorescent protein leading to a decrease in fluorescence. Importantly, mVenus fluorescence is internally normalized by co-expression of mCerulean under a constitutive promoter, allowing comparison between different strains and conditions. At neutral pH c-di-AMP levels increase during growth of the WT and Δ*cdaR* mutant, with an apparently slightly higher c-di-AMP concentration in the mutant during the entire growth (Figure 1D). At pH 4.5 the c-di-AMP concentration increases in the WT as growth commences. In contrast, the non-growing mutant cells show little c-di-AMP (Figure 1D). This led us to the question of whether growth can be recovered once the Δ*cdaR* mutant is exposed again to neutral pH after incubation at low pH. Indeed, after overnight growth at pH 4.5, the Δ*cdaR* mutant is able to recommence growth when transferred to neutral pH, but is still unable to grow at pH 4.5 (Fig. S1C). In conclusion, CdaR is dispensable at neutral pH, but essential for *B. subtilis* growth in low pH conditions.

### Low pH results in a conformational change in the YbbR domains

Our *in vivo* results show that CdaR has a role in low-pH adaptation, suggesting that CdaR might act as a pH sensor. We hypothesized that the YbbR domains are involved in this sensing. To test this hypothesis, we first checked if pH has an effect on the oligomerization state of the YbbR domains. We performed SEC-MALS experiments on the CdaR protein lacking the transmembrane region and the 8-residue intracellular N-terminus (CdaRΔTM) as well as several shortened constructs (Fig. S1D). We opted for these constructs to facilitate handling, particularly protein solubility, and data interpretation. Our experiments showed that all CdaR constructs exist exclusively as monomers and that pH has no effect on the oligomerization state of CdaR, independent of the number of YbbR domains or the presence of the IDR (Fig. S2).

We also investigated whether low pH could promote unfolding of the CdaR domains – a mechanism reported earlier for the acid stress proteins HdeA/B in *Escherichia coli*(*36*). Far UV circular dichroism (CD) spectra of CdaRΔTM at different pHs show a predominantly β-sheet profile with no signs of unfolding even at a pH as low as 3.5 or a temperature as high as 35°C (Fig. S3A/B). As the acid sensing mechanism of CdaR does not seem to involve oligomerization or unfolding, we wanted to test if a conformational change occurs upon pH change. Tryptophan residues are ideal probes for conformational changes as their fluorescence emission spectra are highly sensitive to the surrounding environment. CdaRΔTM carries a single tryptophan residue in the fourth YbbR domain (Figure 2A), making it an excellent reporter of conformational changes. Indeed, the emission spectrum of tryptophan is unchanged at a pH range from 7.5 to 5.5, but a blue shift of approximately 5 nm occurs once a pH below 5 is reached (Figure 2B and Fig. S3C). The blue shift is not affected by changes in ionic strength (addition of up to 1 M NaCl, Fig. S3D) or by temperature (Fig. S3E). The pH effect is distinct from an unfolding event: adding 8 M urea leads to a red shift, indicating an increased exposure of the tryptophan to solvent (Fig. S3F). Importantly, the pH-induced blue shift is reverted once the buffer is exchanged to pH 7.5 (Figure 2C), demonstrating that the conformational change is reversible. The blue shift is unchanged upon removal of the C-terminal IDR (Figure 2B, middle), indicating that the conformational change occurs within the YbbR domains and not in the IDR. We wanted to find out if the conformational shift happens within a single domain or if it involves inter-domain rearrangements. A construct of YbbR domain 4 alone – the domain that harbors the tryptophan residue – does not show the pH-induced blue shift, showing that the conformational change is not just a local effect. Additionally, the tryptophan spectrum in the single domain construct is red shifted (*λ*max ∼330 nm), indicating higher solvent exposure than constructs with more than one YbbR domain. (Figure 2B, right).

**Figure 2.**
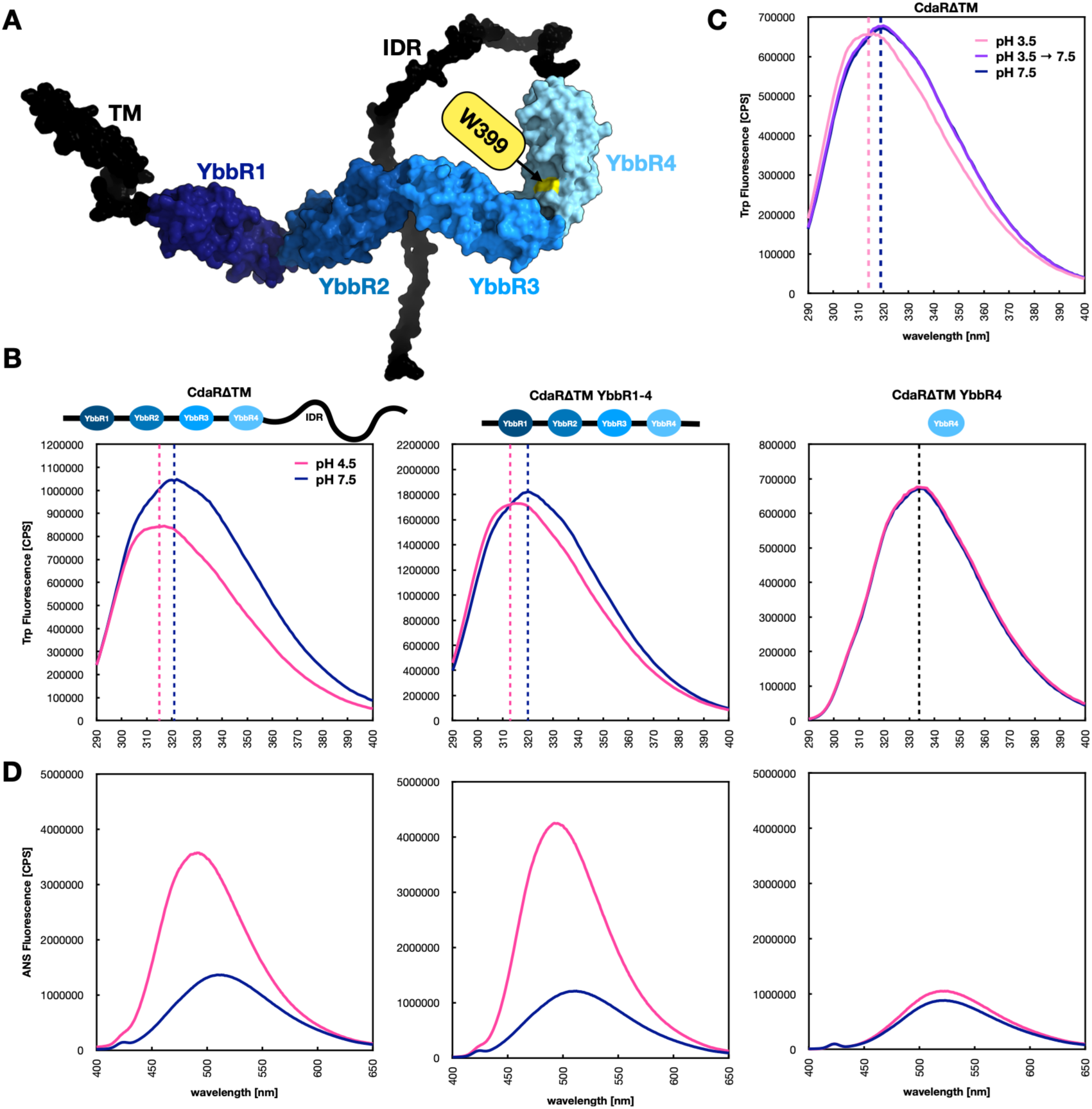
Low pH results in a conformational change in the YbbR domains. A) AlphaFold model of *B. subtilis* CdaR showing the different YbbR domains in shades of blue, the TM and IDR in black and the single tryptophan residue in YbbR domain 4 in yellow. B) Tryptophan fluorescence emission spectra of CdaRΔTM (left), CdaRΔTM YbbR1-4 (middle) and CdaRΔTM YbbR4 (right) at pH 7.5 and 4.5. The two reference lines depict the maxima at pH 4.5 and 7.5. C) Tryptophan fluorescence emission spectra of CdaRΔTM at pH 7.5 (blue), 3.5 (pink) and 7.5 after exposure to pH 3.5 (purple). D) ANS fluorescence emission spectra of the same constructs at pH 7.5 and 4.5.

To narrow down our search for the key region necessary for pH sensing, we used ANS (8-anilinonaphthalene-1-sulfonic acid) fluorescence experiments as the tryptophan fluorescence is limited to constructs containing YbbR domain 4. ANS is commonly used as a fluorescent molecular probe for ligand binding or conformational changes, due to an increase in fluorescence and blue shift in nonpolar environments. Interestingly, all constructs containing two or more YbbR domains show an increase in fluorescence when shifted from pH 7.5 to 4.5, while the single domains show no change in fluorescence (Figure 2D and Fig. S4A). A more detailed titration with CdaRΔTM shows that the change monitored by ANS takes place in the pH range 4.5-5.5 (Fig. S4B/C), similar to what we have observed using Trp fluorescence (Fig. S4D), supporting the idea that both experimental approaches are reporting the same conformational change. Importantly, these experiments show that a minimum of two YbbR domains is necessary to observe a conformational change.

### The hinge region between YbbR domain 1 and 2 is pH-sensitive

In order to better understand the pH adaptation at a molecular level, we decided to determine the structure of the CdaRΔTM protein by X-ray crystallography. However, crystallization trials with CdaRΔTM 1-4YbbR gave no hits. Nevertheless, we obtained crystals from CdaRΔTM 1-2YbbR in many conditions. As we have previously shown that two domains are the minimum pH-responsive unit, we continued our experiments with this construct. Even though we could never grow crystals at a pH below 5, we obtained several structures in a pH range from 5 - 8.5 with resolutions varying from 1.1 to 1.8 Å. As the structures are highly similar to each other and to the one previously solved (PDB: 4QDY) from *S. pneumoniae*, apart from a slight change in the angle between the two YbbR domains, we decided to refine one representative structure at low pH (PDB: 32RB) and one at high pH (PDB: 32QW). Each asymmetric unit contains two chains of CdaRΔTM 1-2YbbR. An overlay of the two chains shows that chain A and B differ only minimally in the angle between YbbR domain 1 and 2 (Fig. S5A). Detailed comparison of the structures determined at different pHs show that the conformation of chain B is independent of pH. In contrast, chain A adopts distinct side-chain conformations in the hinge region connecting YbbR domains 1 and 2 at low and high pH (Figure 3A/B and Fig. S5B). At pH 7.5, R138 is observed in two alternative conformations, oriented either down towards D63 or up towards T174. In the pH 5.0 structure, R138 instead adopts a single upward-facing conformation, forming an interaction with T174. A second pH-dependent rearrangement involves Y66 and K105: at pH 7.5, Y66 contacts K105 from the neighboring loop, whereas in the pH 5.0 structure the two residues are further apart. Finally, H103 interacts with E102 at pH 7.5, while at pH 5.0 E102 adopts two alternative conformations that disrupt this interaction and instead position the side chain toward either D100 or D62 in the adjacent loop.

**Figure 3.**
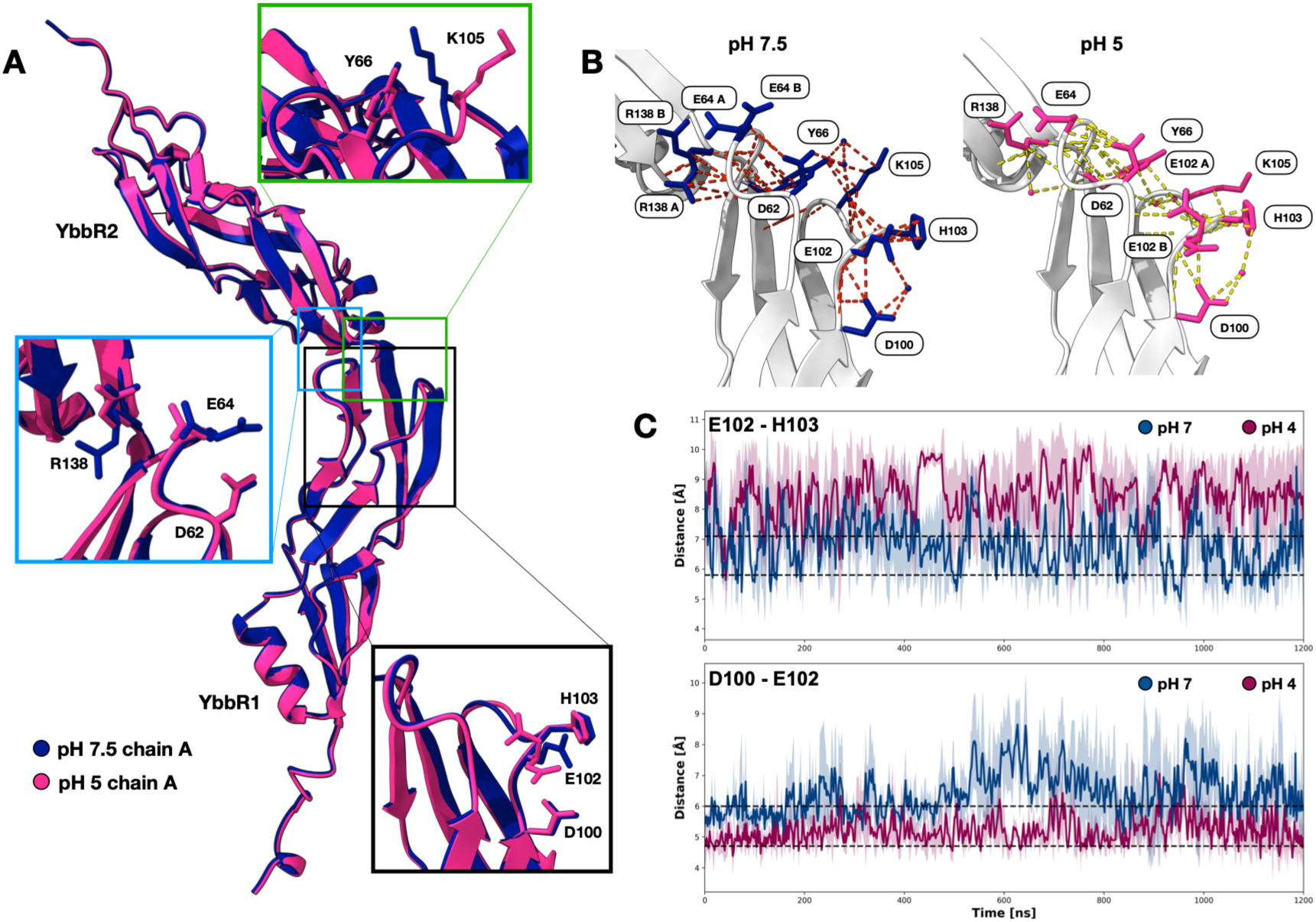
Crystal structures of the YbbR domains at low and high pH show distinct side-chain arrangements. A) Crystal structures of CdaRΔTM YbbR1-2 at two different pHs. In blue pH 7.5 (PDB 32QW), in pink pH 5.0 (PDB 32RB). The three regions that present different side chain arrangement at the two pHs are shown in insets in more detail. **B)** Change of the interaction networks of the residues in the hinge region at pH 7.5 (left, blue) and pH 5 (right, pink). Interactions are shown using the ChimeraX contacts tool with VDW overlap ≥ −0.4 Å. **C)** Distances between the side chains of residues E102 - H103 and D100 - E102 during molecular dynamics simulations. The mean of the three replicate runs is shown as thick lines (dark blue pH 7.0 and purple pH 4.0) and the standard deviation as light pink area (pH 4.0) and light blue area (pH 7.0). The experimentally measured distances between the residues in the X-ray structure at pH 7.5 and pH 5.0 are shown as a black dashed line.

Since using X-ray crystallography, we could not gather experimental data at a pH below 5, we employed molecular dynamics (MD) simulations to gain insights into the interactions mediated by these residues at lower pH. We performed three MD simulations for just over 1 µs for each charge state corresponding to pH 7 and pH 4. Calculating the distance between the center of mass of the side chains of D100, E102 and H103, we can observe the same behavior in MD as we did in X-ray: At low pH residue E102 is closer to D100 and at high pH residue E102 is closer to H103 (Figure 3C).

Contrary to X-ray crystallography, solution-state nuclear magnetic resonance (NMR) is ideally suited to study dynamic systems. 2D ^1^H - ^15^N NMR spectra of ^15^N,^13^C labeled CdaRΔTM 1-2YbbR at pH 7.5 and pH 4.5 exhibited excellent chemical shift dispersion, characteristic of a well-folded protein with a defined tertiary structure at both pHs with distinct chemical-shift changes between the two states (Figure 4A), again excluding unfolding at low pH. In order to gain residue-resolved insights into the conformation and the conformational changes, we used triple-resonance experiments to assign the backbone resonances to a very large extent (93% at pH 4.5, 80% at pH 7.5, Fig. S6A). The chemical-shift perturbations (CSP) observed between the pH 7.5 and 4.5 spectra are localized in the hinge region between the two YbbR domains (Figure 4B/C), with the largest effects observed in the loop comprising residues 100-109. Of note, low peak intensities around residue 100 point to millisecond dynamics of this hinge (Fig. S6B); residue 102 was not observable in assignment experiments, presumably due to these exchange processes. The observed chemical-shift pattern is not influenced by the addition of salt or the use of different buffers (Fig. S6C/D/E), showing that the observed pH effect in this region occurs independently of the exact conditions. Several charged residues are found in this and the adjacent loop (D62, D63, E64, D100, E102, H103 and K105) (Figure 4D). As typical side-chain pK_a_ values of several of these residues fall into the pH region at which the conformational transition is observed, the change of their protonation state provides a plausible mechanism for the observed conformational changes. Altogether, the crystal structure, MD and NMR experiments strongly indicate that the hinge region between YbbR domains is a focal point in the pH-dependent conformational change.

**Figure 4.**
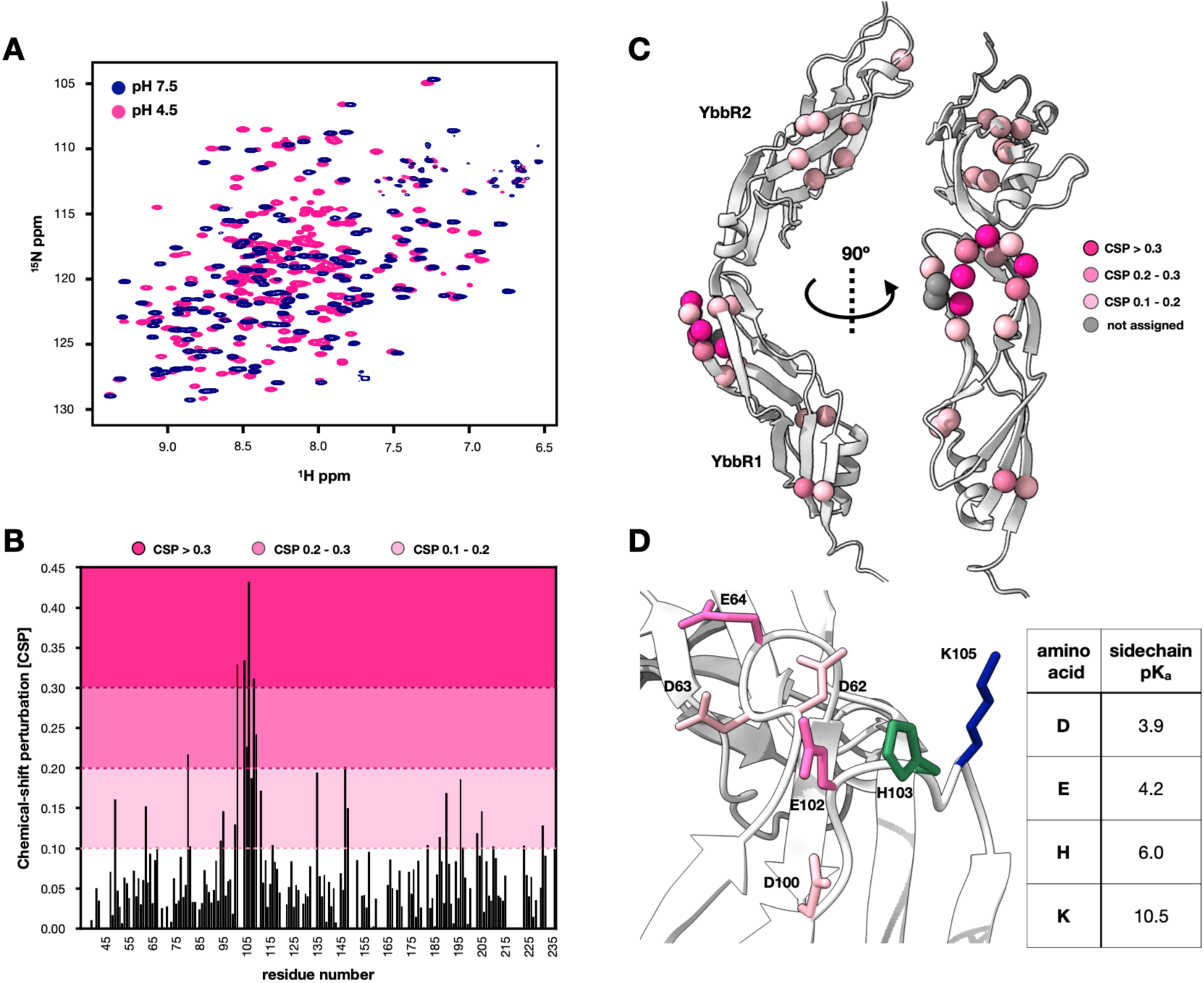
The hinge region between YbbR domain 1 and 2 is pH-sensitive. A) ^1^H-^15^N HSQC spectra of CdaRΔTM YbbR1-2 at pH 7.5 (blue) and pH 4.5 (pink). B) The chemical-shift perturbation (CSP) between the two spectra at pH 7.5 and 4.5 is shown per residue. C) The CSP plotted on the crystal structure of CdaRΔTM YbbR1-2. The colour code and the used cut-off are shown on the righthand side. Of note the hinge region between domain 1 and 2 exhibits the largest CSP. D) Charged residues in the hinge region, which shows the largest chemical-shift changes, colored according to their pK_a_.

### Low pH stabilizes the hinge region and reduces YbbR domain motion

A more detailed analysis of the MD simulation at low and high pH (Figure 5A) reveals global structural changes during the entire MD simulation. Three regions in particular show large differences in their secondary structure between low and high pH (Figure 5A, lower panel): (i) in region 3, a helix forms at low pH around residue 155 (Figure 5B, top and Movie S1), while it is in a disordered state at high pH; (ii) regions 1 and 2, comprising residues 58-62 and 100-104 adopt a coil conformation at high pH and fold into a beta-strand or beta-turn at low pH (Figure 5B bottom and Movie S2/S3). These two last segments form the hinge region discussed above. This is supported by our experimental NMR data that report equally on the residue-wise secondary structure (Figure 5C, top) and the backbone N-H order parameters (bottom): these data show that at pH 4.5 a helix is formed around residue 155, and that the region from residue 100-104 is highly ordered. On the residue level, the MD simulation of the two states shows that at high pH, when the hinge region is unfolded, the conserved D100 is interacting with K114 and not with the residues of the hinge region (Figure 5B and 5D). However, at low pH, D100 is turned towards the hinge region, interacting with E102 as well as with residues from the adjacent loop (residue 60-62).

**Figure 5.**
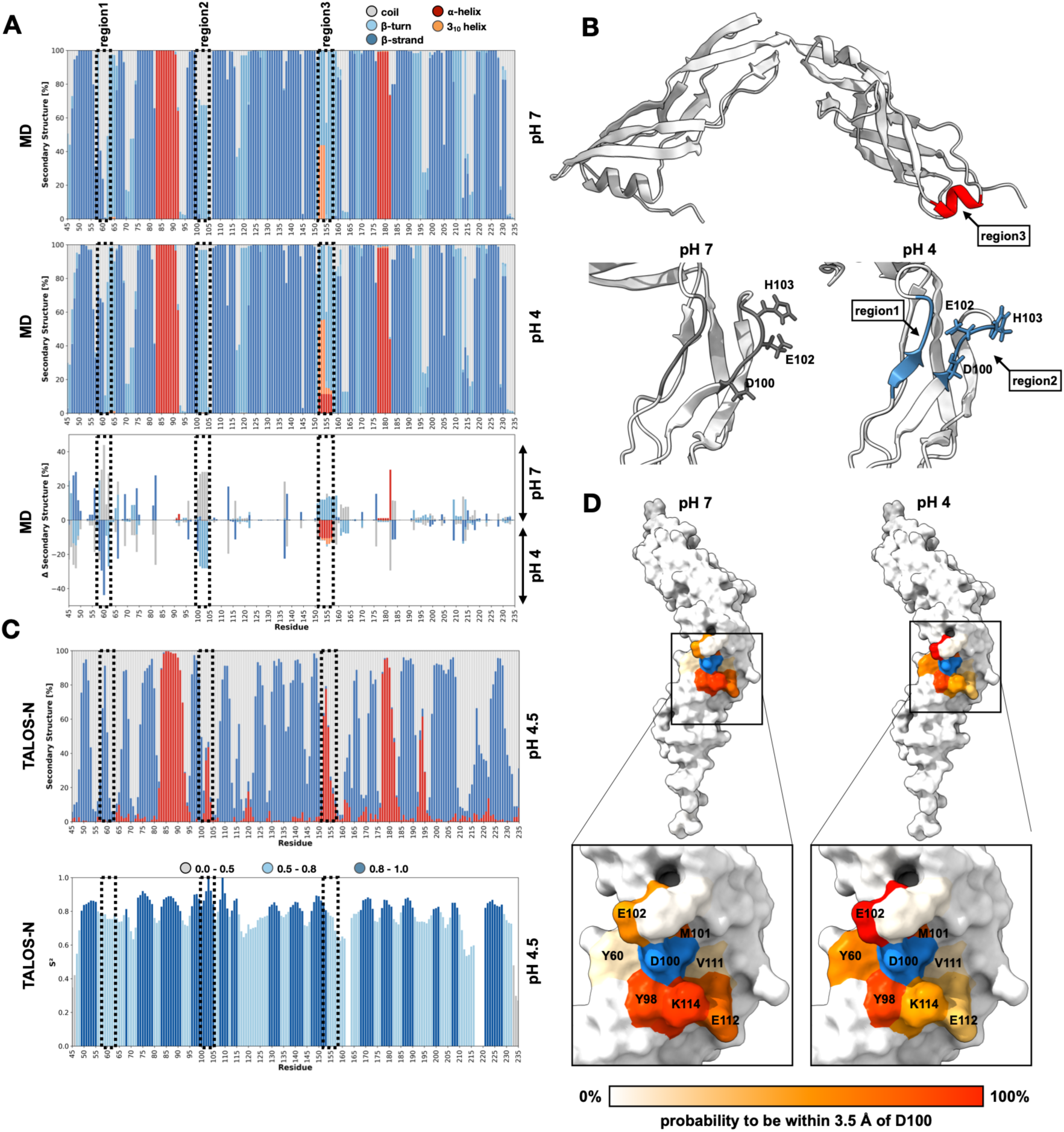
Low pH stabilizes the hinge region. A) Residue plots of CdaRΔTM YbbR1-2 protein showing the probability of having a certain secondary structure during the MD simulation. Top: pH 7.0, middle: pH 4.0 and bottom: differences in secondary structure elements between the two conditions. The regions showing the highest secondary structure changes are highlighted in all graphs with dashed lines, excluding the highly flexible N-terminus. B) Examples of the different secondary structures formed at low pH. Top: increased alpha helical propensity of residues 152-157 shown at pH 4.0 (run3 frame 654), bottom: the hinge region at pH 7 (run1 frame 300) showing less interaction of residues 58-62 with residues 100-104 than at pH 4 (run3 frame 300). C) Secondary structure (top) and order parameter (S2, bottom) prediction of the CdaRΔTM YbbR1-2 protein based on the chemical-shift assignment at pH 4.5 using the software TALOS-N. D) The probability to be within 3.5 Å of residue D100 during the three MD simulations either at pH 7 (left) or pH 4 (right). D100 is colored in blue and the residues in close contact in shades of red, depending on their occupancy over the entire simulation.

All data above have pointed to a change in the hinge region as the key pH sensor. We wondered if this mechanism involves a change in the relative orientation of the YbbR domains – which are connected by this linker. The NMR spectrum of the construct comprising two YbbR domains is, to a very good approximation, the sum of the spectra of the individual domains (CdaRΔTM 1YbbR and CdaRΔTM 2YbbR – Fig. S7A), indicating that there are no extensive contacts between the domains. This can be also appreciated from the MD data: aligning YbbR1 along the trajectories shows that YbbR2 can adopt nearly any conformation during the simulation (see Fig. S7B).

To dissect the different movements, we analyzed the torsion angle between the two domains and the distance of the center of masses of YbbR1 and YbbR2 and noted differences between the two pH conditions. In particular, we observed large changes in the torsion angle at high pH but not at low pH. The torsion angle was as high as 180° in run1 at pH 7, leading to a “flip” of YbbR1 with respect to YbbR2 (Fig. S7C and Movie S4). This flip is synchronized with unfolding of the hinge region, which can only be observed in the high pH simulations. In contrast, at low pH we observed an increase in RMSD and a decrease in inter-domain distance and torsion angle at around 900 ns that corresponds to a “closing” of the two domains (Fig. S7D and Movie S5).

Collectively, our data revealed a pH-dependent conformational switch occurring in the hinge region between YbbR domain 1 and 2, which leads to a rigidification of the hinge region and a subsequent restriction of interdomain dynamics.

### Biochemical and functional relevance of the hinge region

We wanted to know if the residues identified in our structural studies are key to observing a conformational change at low pH using tryptophan fluorescence. To this end, we created individual mutants of the charged residues in the affected hinge region that show the highest chemical-shift perturbation (D62, D100, E102, H103, K105) in CdaRΔTM. We performed tryptophan fluorescence experiments with all protein mutants and discovered that none of the mutants is able to undergo the conformational change (Figure 6A), demonstrating that the residues identified with the construct YbbR1-YbbR2 are responsible for the conformational change observed in the construct with the full-complement of YbbR domains. In addition, these experiments also suggest that the whole network of charged residues in the hinge region participates in the pH switching mechanism.

**Figure 6.**
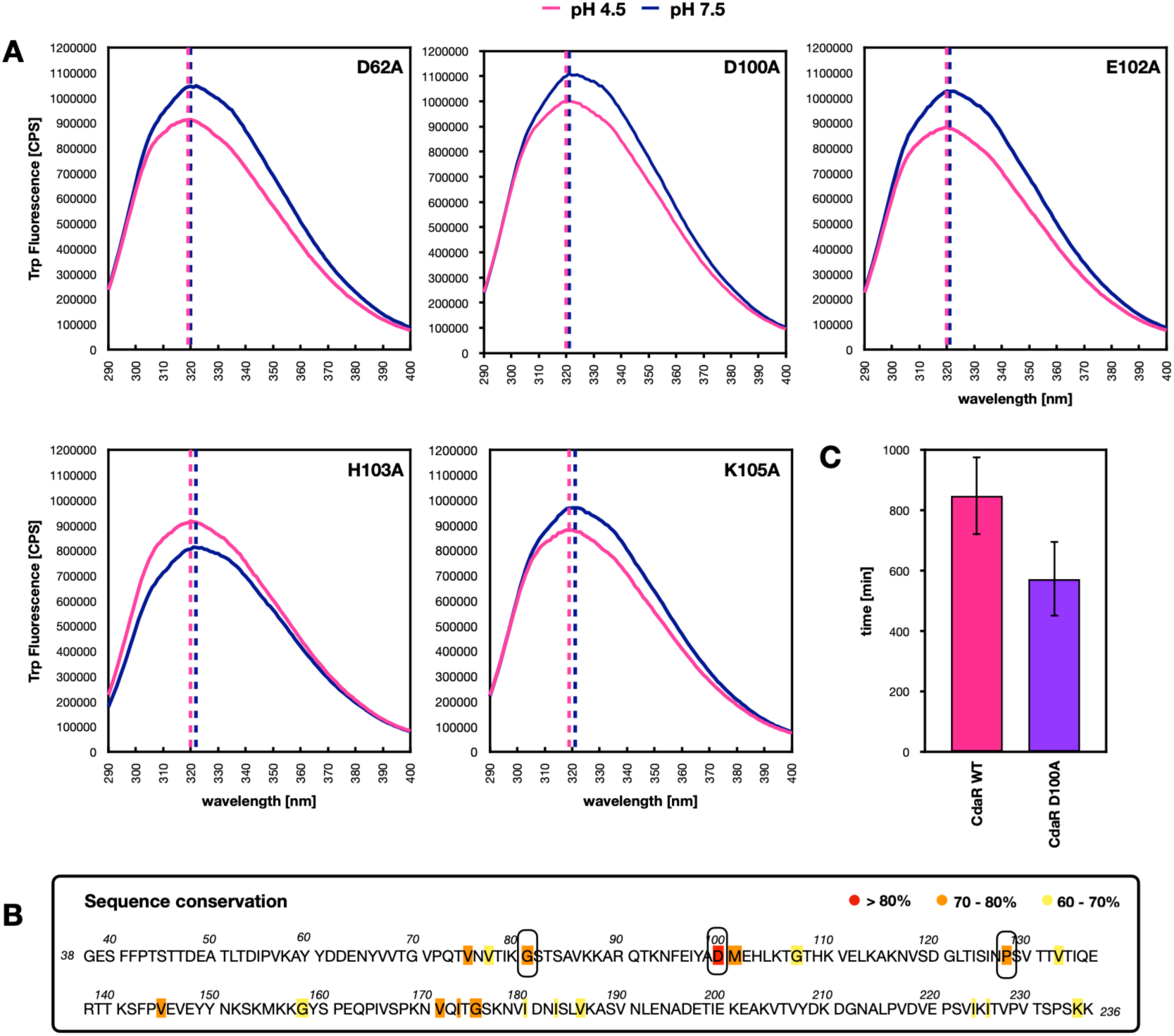
A H-bonding network of charged residues centered around the conserved D100 is responsible for pH sensing. A) Tryptophan fluorescence emission spectra of CdaRΔTM mutants at pH 4.5 (pink) and pH 7.5 (blue). B) Conservation mapped onto the *B. subtilis* CdaR sequence, derived from a ClustalΩ multiple sequence alignment of YbbR domain containing proteins deposited in the AlphaFold database. C) Onset of growth in LB_BS_ pH 4.5 of *B. subtilis* WT and a CdaR D100A mutant, calculated as the point when the initial cell density has doubled (WT n=8, D100A n=5, error bars show SD).

We wondered if any of the residues in the pH-sensitive patch is conserved across YbbR domains. Multiple sequence alignment using >14 000 protein sequences containing predicted YbbR domains (of different length and with or without diadenylate cyclase activity) from the UniProt database reveals that only residues G81, D100 and P128 are highly conserved across all YbbR proteins (Fig. S8A). As there might be wrongly annotated YbbR domains in the UniProt database, we also tried a structure-based search using the AlphaFold database containing 1229 proteins. Similarly, D100 turns out to be the most conserved residue in the extracellular part of CdaR, with a conservation level of more than 80% (Figure 6B). Additionally, the evolutionary conservation profile of *B. subtilis* CdaR provided by the ConSurf database(37) (Fig. S8B) shows that the hinge region between YbbR domain 1 and 2 is the most conserved part of the protein (besides the TM region). Altogether, this conservation strongly suggests a functional relevance for this residue.

Finally, we analyzed the functional role of D100 in *B. subtilis* by generating a mutant strain where the D100A mutation was introduced in CdaR. This mutant strain shows an earlier onset of growth at pH 4.5 compared to the WT (Figure 6C, Fig. S8C), indicating that it adapts faster to low pH conditions. Although this result appears counterintuitive, it demonstrates that D100 has a physiological role in pH adaptation. The faster adaptation to low pH can be understood if the alanine substitution abolishes the pH-dependent conformational change (Figure 6A) by stabilizing a low-pH conformation of the hinge region.

Collectively, these data identify the loop region between YbbR domain 1 and 2, and particularly the highly conserved D100, as the extracellular pH sensor of the CdaA-CdaR complex.

## Discussion

Unicellular organisms must rely on specific molecular mechanisms to sense and respond to environmental changes. However, the inherent complexity of biological systems often makes it challenging to disentangle and understand these processes. The protein complex formed by CdaR and CdaA represents an example where structural information remains limited and *in vitro* models are technically demanding and often fail to recapitulate *in vivo* activity. Studying the individual partners of this system is therefore a valuable strategy, as it allows tighter control of experimental conditions and can provide mechanistic insights that are otherwise hard to grasp using the full complex. In this study, we demonstrated that CdaR plays a critical role in supporting cell survival under acidic stress in *B. subtilis*, particularly below pH 5. Cells lacking CdaR are unable to divide at pH 4.5, but restart growth once they are transferred to neutral pH medium, showing that CdaR helps *B. subtilis* to thrive in low-pH conditions.

We propose that the sensing mechanism does not involve oligomerization or unfolding of the YbbR domains but a conformational change that occurs once a pH below 5 is reached. We could show that a minimum of two YbbR domains is necessary to observe the conformational change. This is in line with the observation that only ∼2 % of YbbR-domain-containing proteins in the AlphaFold database are shorter than 200 residues (38) (the length expected for a CdaR protein with 1 YbbR domain), several of which do not adopt a typical YbbR domain fold. Additionally, multiple sequence alignment using YbbR sequences annotated in the UniProt database(39) with less than 200 residues results in very poor sequence alignment, suggesting that these proteins are either wrongly annotated or perform functions different from the YbbR domains of the CdaR protein. Furthermore, we were able to pinpoint the pH sensor to a conserved hinge region located between YbbR domain 1 and 2 composed by multiple charged residues that undergo a local pH-dependent conformational change that is propagated to the YbbR1-YbbR2 module by reducing its dynamics at low pH.

Importantly, the pH effect is not generalized to all charged residues, as the YbbR domains carry many Asp/Glu residues equally distributed over the protein and not concentrated in the hinge region (Fig. S9A). Other environmental factors such as salt concentration or temperature had little effect on the structure of the YbbR domains. We speculated that charged amino acids with a pK_a_ around pH 4, such as Asp and Glu residues, might be involved in the sensing mechanism. The hinge region between YbbR domain 1 and 2 contains two possible candidates: D100 and E102. While E102 is not conserved among YbbR domains of different species, D100 is highly conserved, being the most conserved residue in YbbR domains deposited in the AlphaFold database. We confirmed by tryptophan fluorescence that a D100A mutant of CdaRΔTM shows no blue shift at low pH *in vitro* and therefore no conformational change. Importantly, this mutant shows a clear change in adaptation to low pH *in vivo* compared to the WT (Figure 6C), adapting faster. The explanation for this effect is not straightforward, one hypothesis might be that Ala more closely resembles a protonated uncharged Asp than to the deprotonated form. We interpret this as evidence that D100A locks the hinge in a conformation resembling the low-pH state rather than simply disabling signaling, effectively pre-adapting the protein. The exact molecular mechanism of pH sensing is not completely clear. A possible trigger for the conformational change might be the opening of a salt bridge at the target pH. However, in our MD simulation, none of the charged residues in the hinge region form a stable salt bridge during the 1 µs simulation. Furthermore, the distance between D100 and the next possible partner K114 is between 5 and 7 Å in the two crystal structures (PDB: 32RB/32QW). Thus, rather than relying on the disruption of a salt bridge, the conformational change might be mediated through an increased H-bonding network, as can be observed in the MD simulation at pH 4 (Fig. S9B), and the exclusion of water (Fig. S9C/D) stabilizing the YbbR domains in a more rigid conformation.

Brogan et al. have recently shown that the IDR of *B. subtilis* CdaR is important for monitoring cell wall damage(33). Nevertheless, the fact that the IDR is not a conserved feature in CdaR proteins, with only about 25-50% (Fig. S9E) of all YbbR containing proteins carrying an IDR, at the C-terminus or in between the YbbR domains, makes it unlikely that peptidoglycan damage sensing is the only purpose of CdaR. The role of the YbbR domains of CdaR has been enigmatic for a long time as the effect on the activity of CdaA is highly context-dependent. It has been shown that CdaR can have both activating and inhibiting effects on CdaA. Similarly, our results demonstrate that while CdaR seems to be dispensable at neutral or alkaline pH for cell survival, showing no effect on CdaA activity, it is essential for cell growth and c-di-AMP production at pH levels below 5. Thus, in pathogens, CdaR might serve as a sensor that is only activated when the bacterium encounters hazardous conditions such as in lysosomes or the stomach. But not only pathogens are faced with variable extracellular pH: during fermentation, pH levels can drop to 3 - 4, and soil bacteria also thrive in natural environments across a wide pH range. Another important aspect is cell survival in bacterial communities. Tran et al. have shown that the pH in *B. subtilis* biofilms drops drastically due to an accumulation of acids within the biofilm(19). Bacteria react with alkalinization once a pH of ∼5.5 is reached. This mechanism of pH adaptation is crucial for cell growth and survival. In this context, we show here for the first time that *B. subtilis* can actively change the external pH to neutral conditions across a wide range of starting-pHs and only resumes growth when reaching favorable pH conditions.

To conclude, we propose a sensing mechanism for CdaR by which pH change is detected by a network of charged residues centered around a conserved Asp residue in the hinge region between YbbR domain 1 and 2, leading to a conformational change that eventually must be transmitted to the intracellular diadenylate cyclase domain of CdaA.

## Materials and Methods

### Cloning

*Bacillus subtilis* CdaR without the predicted transmembrane region (CdaRΔTM - protein sequence 38-484) was PCR amplified from chromosomal DNA and subsequently cloned into a pET28c vector using restriction sites NdeI and XhoI resulting in an N-terminally His-tagged construct with Thrombin cleavage site. The shortened constructs were obtained using CdaRΔTM and site directed mutagenesis introducing Stop codons at protein sequence 417 (CdaRΔTM 1-4YbbR), 326 (CdaRΔTM 1-3YbbR), 237 (CdaRΔTM 1-2YbbR) and 143 (CdaRΔTM 1YbbR). The other single domains were obtained using the above constructs and site directed deletion mutagenesis resulting in the constructs CdaRΔTM 2YbbR (protein sequence 130-236), CdaRΔTM 3YbbR (protein sequence 229-325) and CdaRΔTM 4YbbR (protein sequence 317-416).

### Protein expression and purification

All constructs were expressed in *E. coli* BL21 (DE3). Cells were grown until OD_600_ of 0.7, induced with 0.5 mM IPTG and incubated overnight at 20°C. The cell pellet was resuspended in 30 mL lysis buffer A (50 mM Tris-HCl pH 7.5, 500 mM NaCl, 20 mM Imidazole, 1 mM PMSF), homogenized using an EmulsiFlex homogenizer and the His-tagged protein purified on a 5 mL GE HisTrap FF Ni-Sepharose. The lysate was clarified by centrifugation and the supernatant loaded on the HisTrap. After washing with 10 column volumes (CVs) wash buffer A (50 mM Tris-HCl pH 7.5, 300 mM NaCl, 20 mM Imidazole) the protein was eluted using a step gradient of imidazole (100, 200, 300, 400 and 500 mM 2 CVs each). The eluted protein was dialyzed and Thrombin cleaved overnight against SEC buffer A (50 mM Tris-HCl pH 7.5, 150 mM NaCl) and further purified on a Superdex 200 16/600 column in SEC buffer A.

### *In vivo* experiments

All *in vivo* experiments were performed in *B. subtilis* 168 strain (WT). The Δ*cdaR* mutant was generated using a two-step mutagenesis protocol adapted from Koo et al (40). Briefly, the *cdaR* gene was replaced with the kanamycin resistance cassette (BKK) by transforming the WT strain with a PCR fragment containing the BKK cassette flanked by 1 kb of *cdaR* flanking regions. Subsequently, the Δ*cdaR*::BKK strain was transformed with the pDR244 plasmid to excise the antibiotic marker via Cre recombinase. The final markerless Δ*cdaR* mutant, confirmed by PCR and sequencing, contains a 150 bp scar sequence at the *cdaR* locus.

Cells were grown in 10 g/L Tryptone, 5 g/L yeast extract and 5 g/L NaCl (LB_BS_). pH was adjusted using HCl or NaOH. For Agar plates 15 g/L Agar was added to the medium. To measure pH during cell growth 15 mg/L phenol red for liquid cultures and 25 mg/L phenol red for Agar plates were added to the medium. For Agar plates 3 mL cells were grown at 37°C until OD_600_ of 1 was reached. The cell pellet was washed with 150 mM NaCl and a serial dilution (10^-1^ to 10^-6^) made. Of each dilution 5 µl were spotted on the Agar plate and the plate incubated at 37°C. Growth is recorded after 24h.

For liquid cultures cells were grown in LB_BS_ pH 6 at 37°C for 4h. Subsequently the cells were harvested and the cell pellet washed with the same volume LB_BS_ pH 6 as the initial culture. In a 96 well plate cells were inoculated at OD_600_ 0.1 in media with pH 4.5-9, both with and without phenol red as a pH indicator. In a CLARIOStar plate reader an absorption spectrum from 350-750 nm is recorded every 15 min for 16 hours to follow both pH and OD_600_.

### Microscopy

The morphology of *Bacillus subtilis* 168 (wild type) and a Δ*cdaR* mutant strain was examined under neutral and acidic growth conditions. Cells were initially streaked on LB_BS_ agar and grown overnight at 37°C. Pre-cultures were prepared in LB_BS_ medium (10 g/L tryptone, 5 g/L yeast extract, 5 g/L NaCl) adjusted to pH 6.0 and incubated at 37°C with shaking (220 rpm) until an OD600 of approximately 1.0 was reached. Cells were harvested by centrifugation (4000 rpm, 5 min), washed with fresh medium, and diluted to an OD_600_ of 0.1 in LB_BS_ adjusted to pH 7.0 or pH 4.5. Cultures were grown overnight at 37°C with shaking.

For fixation, cultures were mixed 1:1 with a fixative solution containing 4% paraformaldehyde, 0.04% glutaraldehyde, and 200 mM sodium phosphate buffer (pH 7.0). Samples were incubated for 15–30 min at room temperature, followed by 1 h on ice. Fixed cells were pelleted (5000 rpm, 2 min), washed twice with 100 mM sodium phosphate buffer (pH 7.0), and resuspended in the same buffer. Cell densities were adjusted as required prior to imaging.

For microscopy, 2 μL of fixed-cell suspension was deposited onto a glass-bottom µ-Slide 8 Well chamber. Cells were immobilized using a freshly prepared 1% agarose pad and imaged using a Nikon Eclipse Ti microscope equipped with a CFI Plan Apochromat Lambda D 60× oil immersion objective (NA 1.42) and an Andor iXon 888 camera. Images were acquired at an effective magnification of 90× (60× objective with 1.5× intermediate magnification).

### Endospore staining

Endospore formation was assessed in *Bacillus subtilis* 168 (wild type) and a Δ*cdaR* mutant grown under neutral and acidic conditions. Strains were initially streaked on LB_BS_ agar and incubated overnight at 37°C. Pre-cultures were prepared in LB_BS_ medium (10 g/L tryptone, 5 g/L yeast extract, 5 g/L NaCl) adjusted to pH 6.0 and grown at 37°C with shaking (220 rpm) until reaching an OD600 of approximately 1.0. Cells were harvested by centrifugation, washed with fresh medium, and diluted to an OD600 of 0.1 in LB_BS_ adjusted to pH 6.0 or pH 4.5. Cultures were incubated overnight at 37°C with shaking.

Cell densities were normalized prior to staining and serially diluted in 150 mM NaCl. Aliquots (20 μL) were spread onto clean microscope slides, air-dried, and heat-fixed. Endospores were stained using the Schaeffer-Fulton method. Briefly, smears were covered with blotting paper saturated with 0.5% malachite green and steamed for 5 min, with additional stain applied as needed to maintain moisture. Slides were rinsed thoroughly with water and counterstained with 0.5% safranin for 30 s. Excess stain was removed by washing with water, and slides were air-dried.

Stained samples were examined by bright-field microscopy using an Olympus CX31 microscope equipped with a 40×/0.65 objective and a Michrome 5 Pro camera. Endospores stained green, whereas vegetative cells appeared pink to reddish-brown.

### cyclic-di-AMP biosensor

The transcriptional c-di-AMP sensor used in this study (CDAsK) was cloned into the *lacA* integrative plasmid pHT2184(41). CDAsK consists of two transcriptional units separated by a terminator sequence. The first transcriptional unit, which serves as an expression control, contains the fast-folding *mCerulean* under the control of the strong constitutive promoter P_grac210_ from pHT2184 and was cloned using the BamHI and XbaI restriction sites. The second transcriptional unit forms the sensing module and comprises six functional components: an upstream terminator(42), the constitutive promoter Ps1(43), the *kimA* riboswitch, the *mVenusNB* fluorescent reporter fused to an SSRA degradation tag(43) and a downstream terminator. This structure was synthesized and cloned into pHT2184::mCerulean between SalI and EcoRI restriction sites. Based on the initial *in vivo* experiments, two stop codons were introduced by PCR Quickchange upstream of the SSRA tag, thereby increasing reporter fluorescence (Primers listed in the table below).

*Bacillus subtilis* wild-type and Δ*cdaR* were transformed with the plasmid pHT2184::CDAsK linearized with NruI using the Koo transformation protocol(40).

Growth experiments for *B. subtilis* cyclic-di-AMP quantification are performed in SMM minimal medium to reduce background fluorescence.

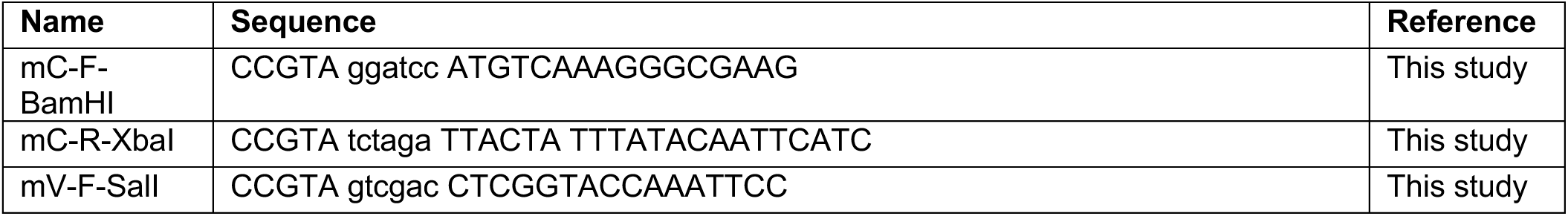
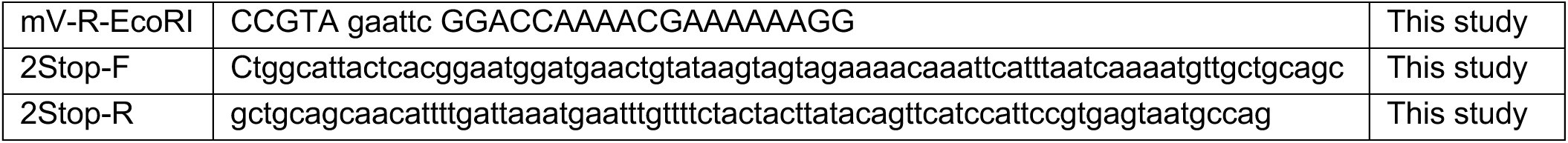

### Size exclusion chromatography coupled to multi-angle light scattering

For determination of particle mass, polydispersity and to study oligomeric state, we used size exclusion chromatography coupled to multiangle-light scattering and differential refractive-index measurements (SEC-MALS). All CdaRΔTM samples were measured in two different buffers (50 mM Tris-HCl pH 7.5, 150 mM NaCl and 50mM sodium acetate pH 4.5, 150 mM NaCl). SEC-MALS runs were performed with 2.0 mg/ml protein concentration for all CdaRΔTM protein samples. Before SEC MALS measurements, all protein samples were centrifuged for 10 min at 16000g at 4°C. 50 - 60 µL of purified protein was injected either onto an SD 200 10/300 Increase column (Cytiva) or onto a Xbridge Premier Protein SEC 250 Å, 2.5 μm column (Waters) connected to a 1260 Infinity II HPLC (Agilent Technologies). A *miniDAWN TREOS II Multi-Angle Light Scattering Detector (*Wyatt Technologies) was used to detect light scattering at 658 nm and an Optilab T-REX (Wyatt Technologies) was used for refractive index measurements. All SEC-MALS runs were performed at room temperature in respective buffer. Average molecular weight was calculated with ASTRA software (Version 8.2.2.119, Water / Wyatt Technology).

### Mass Photometry

Purified CdaRΔTM protein was analyzed by mass photometry in two buffers at pH 7.5 and pH 4.5, corresponding to the buffer conditions used in the SEC-MALS experiments described above. Measurements were done using a Refeyn Two^MP^ mass photometer (Refeyn Ltd., Oxford, UK). Prior to each measurement, the instrument was focused using 10 µL of the respective buffer, after which 1–2 µL of protein sample was added directly to the measurement droplet to achieve a final protein concentration of 25–30 nM. Movies were collected for 60 second and analyzed with DiscoverMP software (Refeyn Ltd.). Instrument calibration was performed in the respective buffer before each data acquisition.

### Fluorescence experiments

For tryptophan fluorescence experiments, all proteins were measured at 2 µM concentration in buffers with different pHs (10 mM sodium acetate pH 3.5/4/4.5/5.5, 10 mM sodium phosphate pH 4.5/5.5/6.5/7.5, 10 mM Tris-HCl pH 7.5 and 10 mM sodium citrate pH 3.5/4.5/5.5/6.5 with or without NaCl 150 mM, 500 mM, 1000 mM). Excitation was performed at 275 nm and the emission spectrum read at 290-400 nm. The same buffers were used for ANS fluorescence measurements. All proteins were measured at a concentration of 0.5 mg/mL in the different pH buffers with 50 µM ANS. Excitation wavelength was 370 nm and the spectrum recorded from 400-650 nm.

### Circular dichroism

Far ultraviolet (FUV) CD spectra of CdaRΔTM were recorded in a 1 mm cell at 0.1 mg/mL protein concentration at 20°C and 35°C in 10 mM sodium acetate pH 3.5/4/4.5/5.5 and 10 mM sodium phosphate pH 4.5/5.5/6.5/7.5 on a Jasco J-815 spectropolarimeter continuously purged with nitrogen.

### X-ray crystallography

CdaRΔTM 1-4YbbR and CdaRΔTM 1-2YbbR were dialyzed into 20 mM Tris-HCl pH 7.5, 150 mM NaCl and concentrated to 10 mg/ml. Initial conditions were screened using commercial crystallization screens with protein concentrations between 5 and 10 mg/mL, a 1:1 protein: reservoir drop ratio, 4 and 20°C in 96 well plates using 45 µl reservoir solution and a drop size of 0.6 µl. While with the CdaRΔTM 1-4YbbR construct no crystals could be obtained, the CdaRΔTM 1-2YbbR crystallized in many conditions which were subsequently optimized. Based on the optimization a final screen was designed containing a pH range from pH 4.5 to 8.5 with different PEG, buffer and salt combinations. Crystals could be obtained at all pHs except below pH 5. Diffraction data were collected on the ALBA BL13-XALOC beamline. Integration was performed on site using the pipeline xia2. Scaling and merging were performed using Aimless included in the software suite CCP4(44). Molecular replacement was done using the software package Phenix(45) and the AlphaFold2 model of *Bacillus subtilis* CdaR without transmembrane region and truncated to YbbR domain 1 and domain 2. Several rounds of phenix.refine were performed and finally the two best structures at pH 5.0 (PDB 32RB) and 7.5 (PDB 32QW) deposited on the wwPDB server.

### Nuclear magnetic resonance

CdaRΔTM, CdaRΔTM 1YbbR, CdaRΔTM 2YbbR and CdaRΔTM 1-2YbbR were expressed in BL21 (DE3) in M9 minimal medium supplemented with ^15^N-ammonium chloride (1 g/L) and or ^13^C-labeled glucose (2 g/L). Expression and protein purification was performed as described above. All NMR experiments were performed on Bruker NEO spectrometers equipped with cryogenically cooled TCI probe heads and operating at magnetic field strengths corresponding to ^1^H Larmor frequencies of 800 and 600, respectively. The sample temperatures were set to 45°, 35° or 25°C in different experiments, as specified. CdaRΔTM 1-2YbbR was assigned in NMR buffer A (20 mM sodium phosphate buffer pH 4.5, 50 mM NaCl, 5% DSS) and in NMR buffer B (20 mM sodium acetate buffer pH 4.5, 50 mM NaCl, 5% DSS) at a protein concentration of 0.36 mM. The following experiments were performed: NMR buffer A 45°C: 2D ^15^N-^1^H BEST TROSY, 2D ^13^C-^1^H HSQC, 3D HNCO, 3D HNCA, 3D HNcaCO, 3D HNcoCA, 3D HNcoCACB, 3D HNCACB, 3D ^15^N-^1^H-^1^H NOESY and in NMR buffer B 35°C: 2D ^15^N-^1^H BEST TROSY, 3D HNCA, 3D HNcoCA, 3D HNcoCACB and 3D HNCACB. Data processing and analysis was performed using the Bruker TopSpin software package and the CCPN software package(46).

### Molecular dynamics simulations

Two systems were prepared from the X-ray structure of CdaRΔTM 1-2YbbR using CHARMM-GUI(47), corresponding to protonation states representative of ∼pH 7 and ∼pH 4. Protonation states of titratable residues were fixed based on average estimated pK_a_ values and manually set-up with CHARMM-GUI PDB manipulation options. At pH 7, Arg and Lys residues were protonated (positively charged), His residues were neutral, and Glu and Asp were deprotonated (negatively charged). At pH 4, Arg, Lys, and His were protonated, while Glu and Asp were neutral, in accordance with predicted pK_a_ values.

The systems were then solvated resulting in a cubic box of dimensions described in the table below. Na^+^ and Cl^-^ions were added using the Autoionize plugin of VMD(48) to neutralize the system resulting in a final concentration of 150 mM NaCl. Three replicates for each system were carried out independently and run until at least 1 µs was simulated.

| **System** | **Protonation state** | **Total Nº of atoms** | **Nº of water molecules** | **Replicate Nº** | **Time [ns]** |
|---|---|---|---|---|---|
| CdaRΔTM 1-2YbbR | pH 7 | 180475 | 59005 | 1 | 1198 |
|  |  |  |  | 2 | 1604 |
|  |  |  |  | 3 | 1191 |
|  | pH 4 | 181110 | 59214 | 1 | 1288 |
|  |  |  |  | 2 | 1072 |
|  |  |  |  | 3 | 1311 |

Simulations were performed with NAMD2(49). The CHARMM36 force field(50) was used to model the protein and ions, and the TIP3P model was chosen for the water. The Particle Mesh Ewald method was used for the treatment of electrostatic interactions, with an upper threshold of 1 Å for grid spacing(51). Electrostatic and van der Waals forces were calculated in every time step with a 12 Å cut-off distance. A switching distance of 10 Å was chosen to smoothly truncate the non-bonded interactions. Only atoms in a Verlet pair list with a 13.5 Å cut-off distance (reassigned every 20 steps) were considered. The SETTLE algorithm was used to constrain all bonds involving hydrogen atoms to allow the use of a 2 fs time step. The Nose-Hoover-Langevin piston method was employed to control the pressure with a 50 fs period, 25 fs damping constant, and a desired value of 1 atmosphere. The system was coupled to a Langevin thermostat to sustain a temperature of 298 K throughout. VMD was used for analysis of the trajectories using home-made python.

## Acknowledgments

We thank Paula Teixeira for providing us with the *B. subtilis ΔcdaR* strain, the Biochemical and Biophysical Technologies and Advances Light Microscopy i3S Scientific Platform for access and support for fluorescence, CD and light microscopy experiments, the Molecular Biology/Protein Services at the Institute of Science and Technology Austria (ISTA) for technical support and assistance with instrument setup and measurements, the NMR facility of ISTA for excellent support and ALBA Synchrotron, and staff, for access to BL13-XALOC beamline and support. This work was funded by National Funds through FCT - Fundação para a Ciência e a Tecnologia, I.P., under the projects 2023.13636.PEX and 2022.03075.PTDC awarded to KW, and by “La Caixa” Foundation under the project LCF/PR/HR22/52420004, awarded to JHM-C. The authors also acknowledge the computational resources provided by the Hyperion supercomputer at the Donostia International Physics Center (DIPC).

## Declaration of generative AI and AI-assisted technologies in the manuscript preparation process

During the preparation of this work, the author(s) used ChatGPT version 5.5 for editing home-made python scripts used for analysis of the MD trajectories in VMD, additionally Claude version 1.52386.0 was used for manuscript proof-reading. The author(s) reviewed and edited the output as needed and take full responsibility for the content of the published article.

