## Supplemental information for "A reversible YbbR-domain hinge switch senses extracellular pH to control bacterial cyclic-di-AMP signaling"

### Supplementary Figures

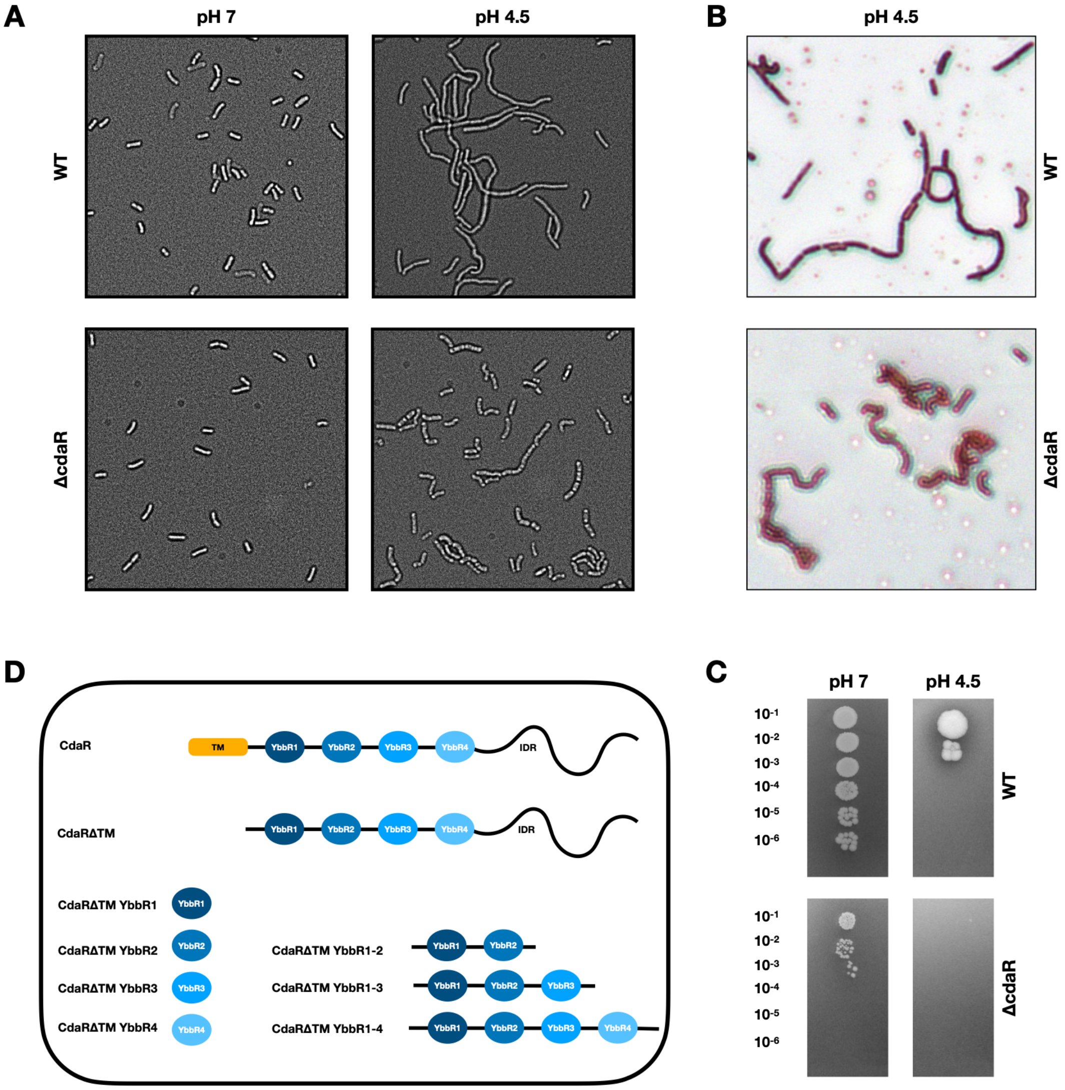


**Fig. S1.**

**A)** Brightfield microscopy image of *Bacillus subtilis* WT and Δ*cdaR* mutant grown in $LB_{BS}$ at pH 7.0 and pH 4.5. **B)** Brightfield microscopy image of *B. subtilis* WT and ΔcdaR in $LB_{BS}$ pH 4.5 stained with safranin and malachite green. **C)** WT and Δ*cdaR B. subtilis* were grown in $LB_{BS}$ pH 4.5 and subsequently plated on $LB_{BS}$ Agar plates with either pH 4.5 or 7. As can be seen on the bottom left image, the Δ*cdaR* mutant can recover growth once transferred from low pH to high pH medium, while it is still unable to grow at pH 4.5 (bottom right). **D)** Depiction of the various CdaR constructs used in this study.

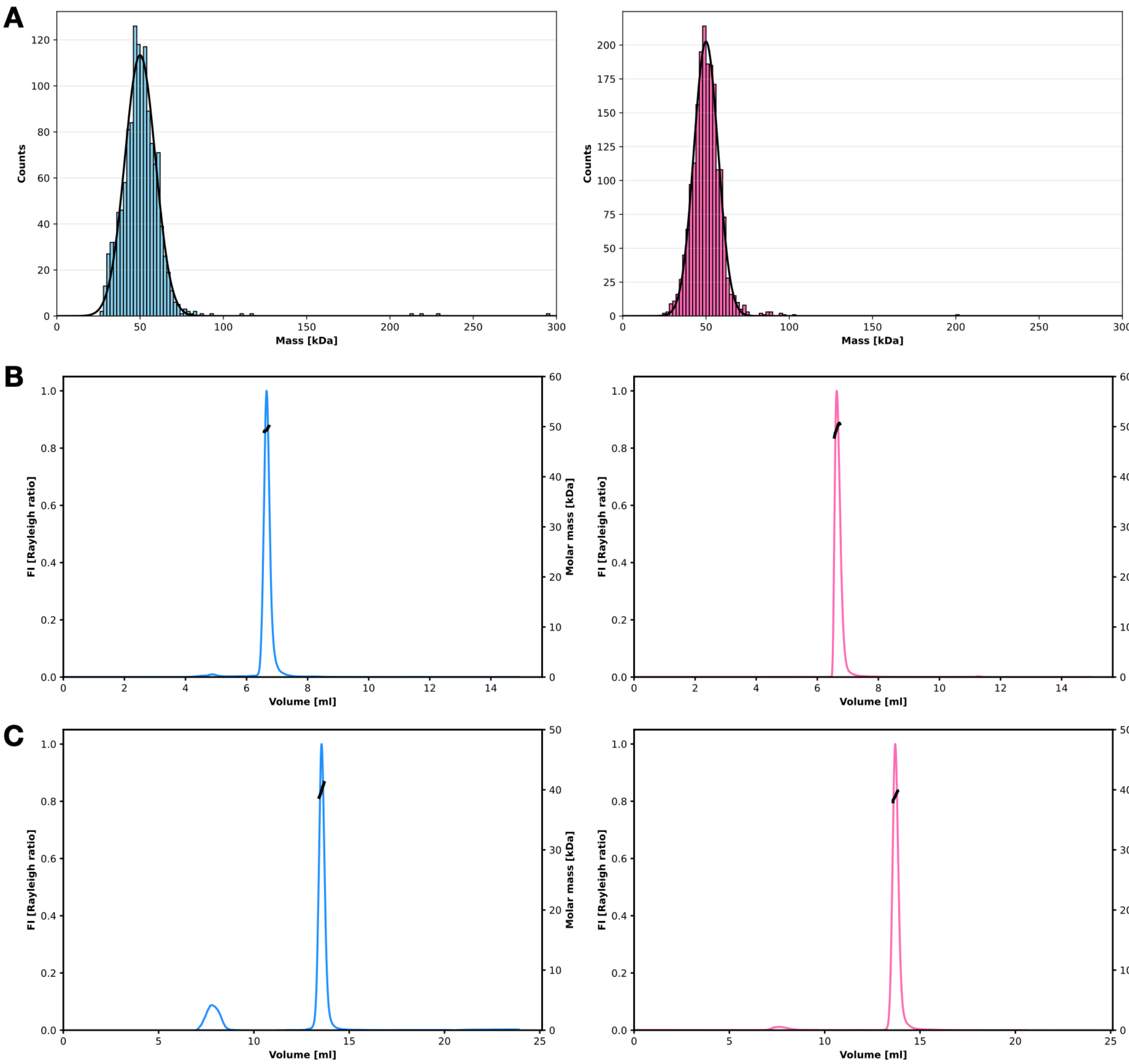


**D**

| construct | estimated MW [kDa] | experimental MW [kDa] | |
|---|---|---|---|
| | | *pH 4.5* | *pH 7.5* |
| CdaRΔTM YbbR1 | 11.90 | 11.91 | 11.90 |
| CdaRΔTM YbbR2 | 12.10 | 12.05 | 12.10 |
| CdaRΔTM YbbR3 | 10.70 | 10.68 | 10.62 |
| CdaRΔTM YbbR4 | 11.30 | 11.50 | 11.43 |
| CdaRΔTM YbbR1-2 | 22.2 | 20.08 | 21.26 |
| CdaRΔTM YbbR1-3 | 31.7 | 29.55 | 30.03 |
| CdaRΔTM YbbR1-4 | 41.7 | 39.96 | 39.98 |
| CdaRΔTM | 50.5 | 49.50 | 49.50 |

**Fig. S2.**

**A)** Mass photometry measurements of CdaRΔTM at pH 7.5 (left, blue) and pH 4.5 (right, pink). Both showing a MW of 50 kDa, the predicted MW of a CdaRΔTM monomer. **B)** SEC MALS profiles of CdaRΔTM at pH 7.5 (left, blue) and pH 4.5 (right, pink). **C)** Same as **B)** but measured on CdaRΔTM YbbR1-4. **D)** Table showing the estimated MW and the experimentally determined MW by SEC MALS of all constructs used in this study. Of note all constructs behave as monomers, both at pH 7.5 and 4.5.

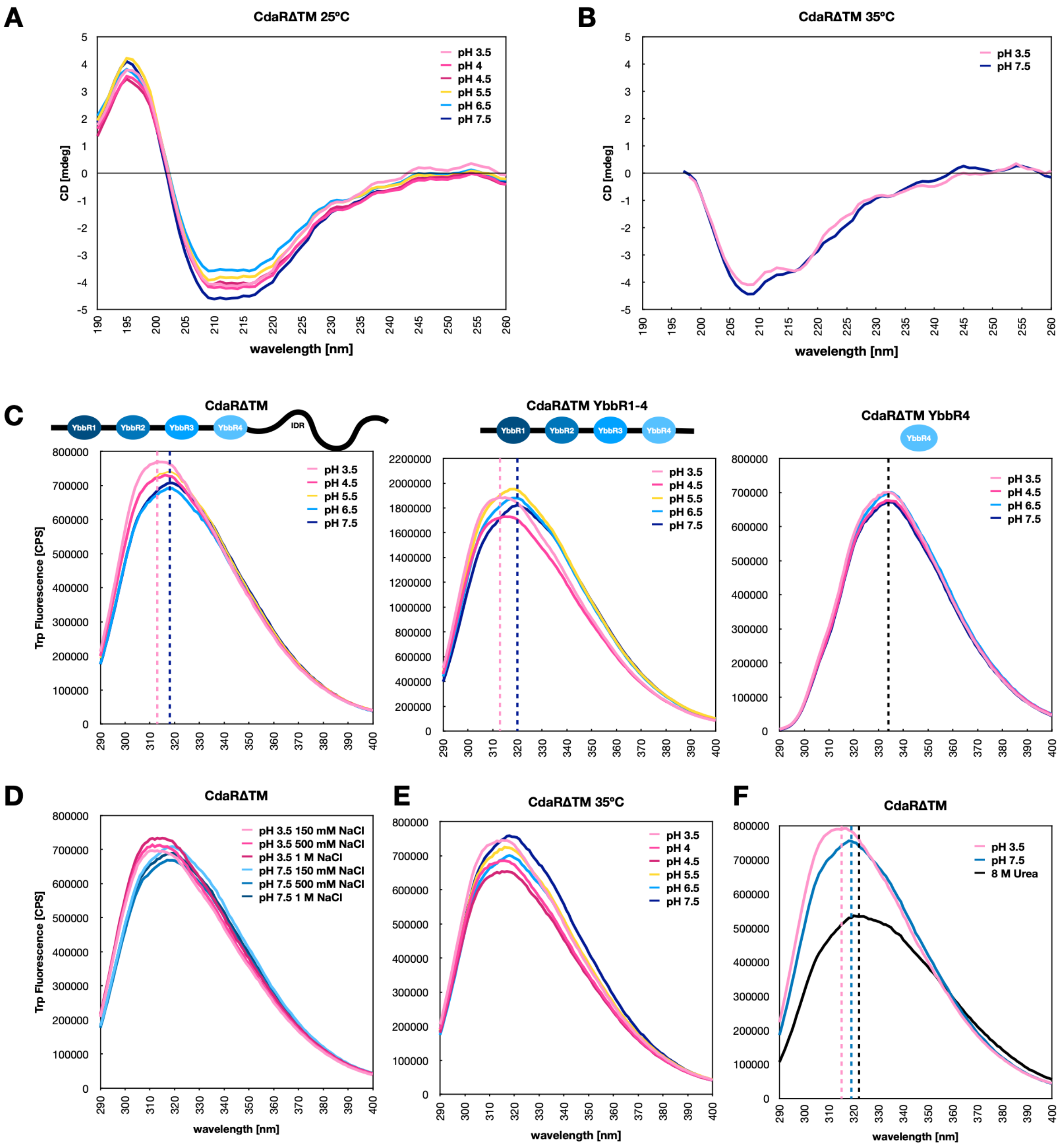


**Fig. S3.**
**A)** Circular dichroism (CD) spectra of CdaRΔTM at different pHs (20ºC). **B)** CD spectra of CdaRΔTM at pH 3.5 and pH 7.5 (35ºC). **C)** Tryptophan fluorescence emission spectra of CdaRΔTM (left), CdaRΔTM YbbR1-4 (middle) and CdaRΔTM YbbR4 (right) at different pHs. **D)** The effect of salt in the tryptophan emission spectra at different pHs. **E)** The same experiment as shown in panel C (right), performed at 35ºC. **F)** Unfolding of the protein with 8 M urea leads to a red shift, contrary to the blue shift induced by exposure to low pH.

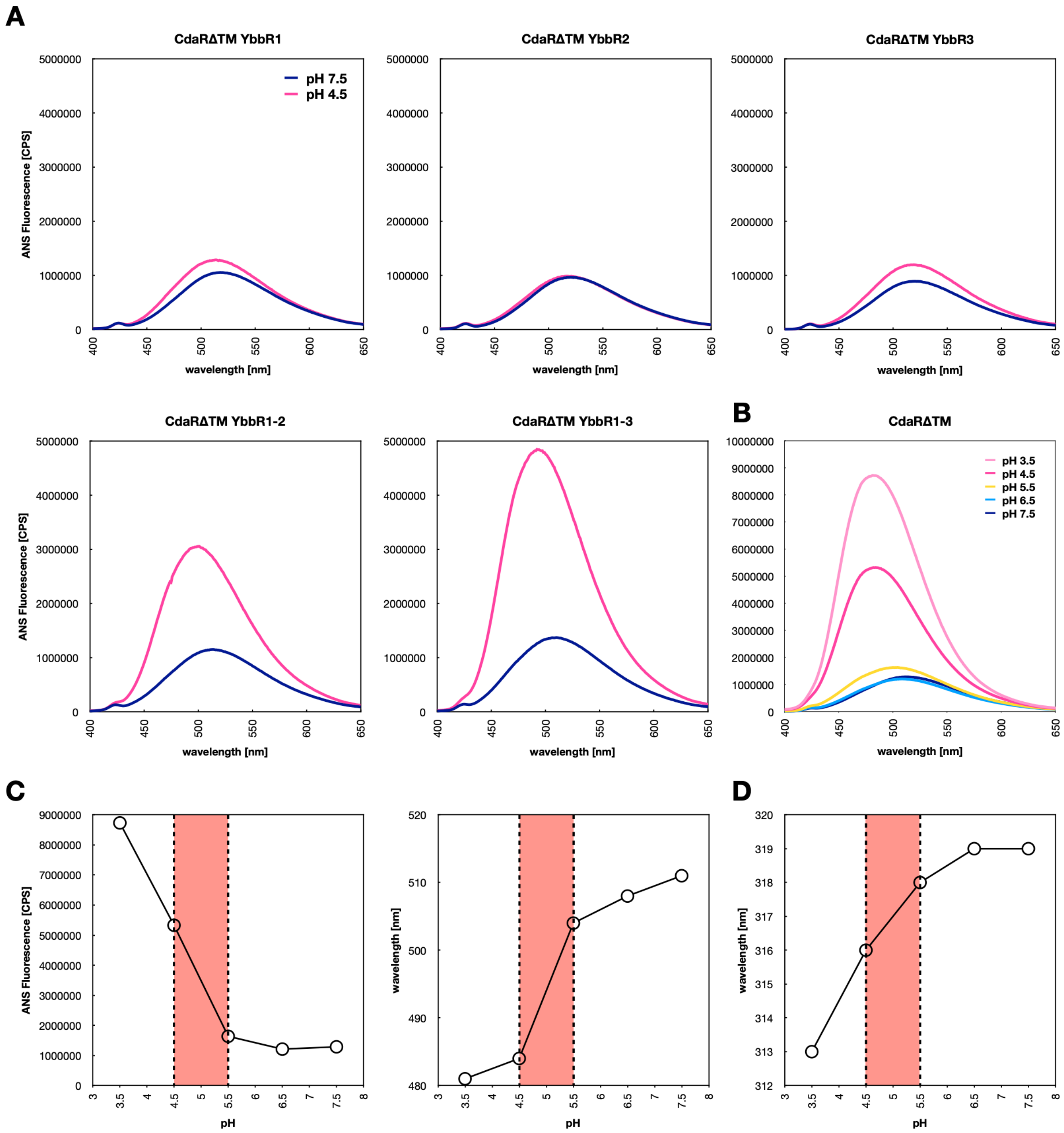


**Fig. S4.**

**A)** ANS fluorescence emission spectra of CdaRΔTM with different numbers of YbbR domains at pH 7.5 and 4.5. **B)** ANS fluorescence emission spectra of CdaRΔTM at different pHs. **C)** pH titration using CdaRΔTM and ANS fluorescence. Both the fluorescence intensity (left) and the wavelength of the peak maximum (right) depend on the pH. In both cases the turning point is between pH 4.5 and 5.5. **D)** pH titration based on the tryptophan fluorescence emission spectra in Supplementary Figure 3C (left), showing that the turning point for the conformational change is between pH 4.5 and 5.5.

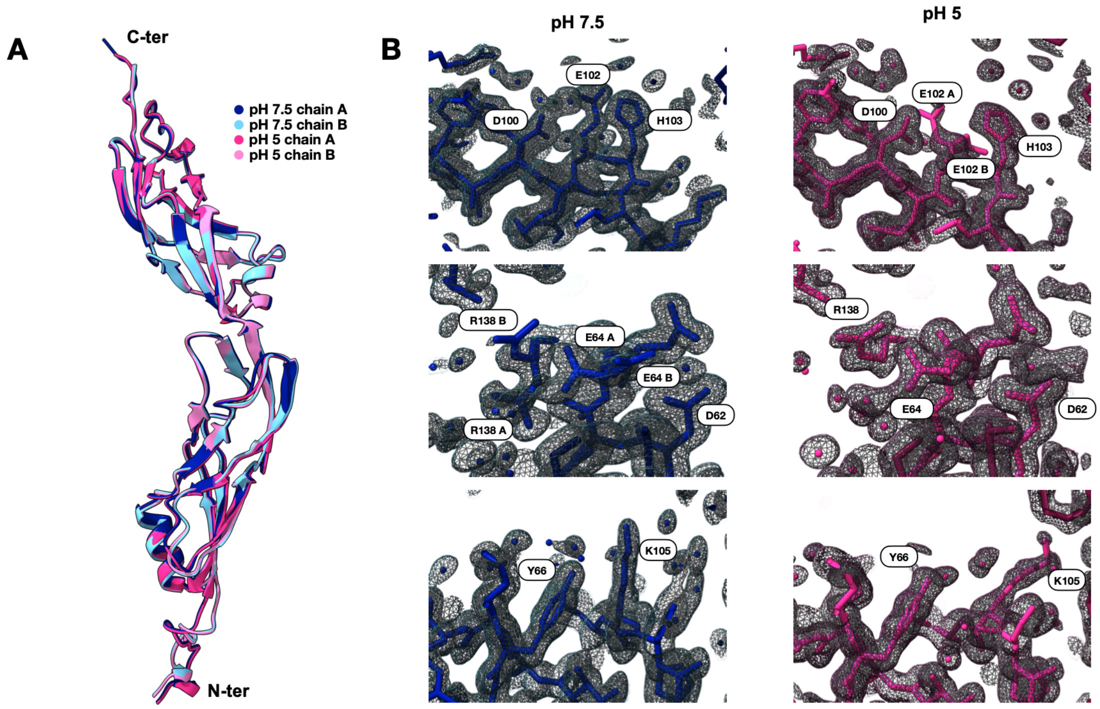


**Fig. S5.**

**A)** Crystal structures of CdaRΔTM YbbR1-2 at pH 7.5 and 5.0 showing only slight differences between chain A and B in the position of the N-terminus and the angle of the two domains with respect to each other. **B)** Regions with distinct side chain arrangement in chain A at pH 7.5 (blue) and 5.0 (pink) with the corresponding 2Fo-Fc map.

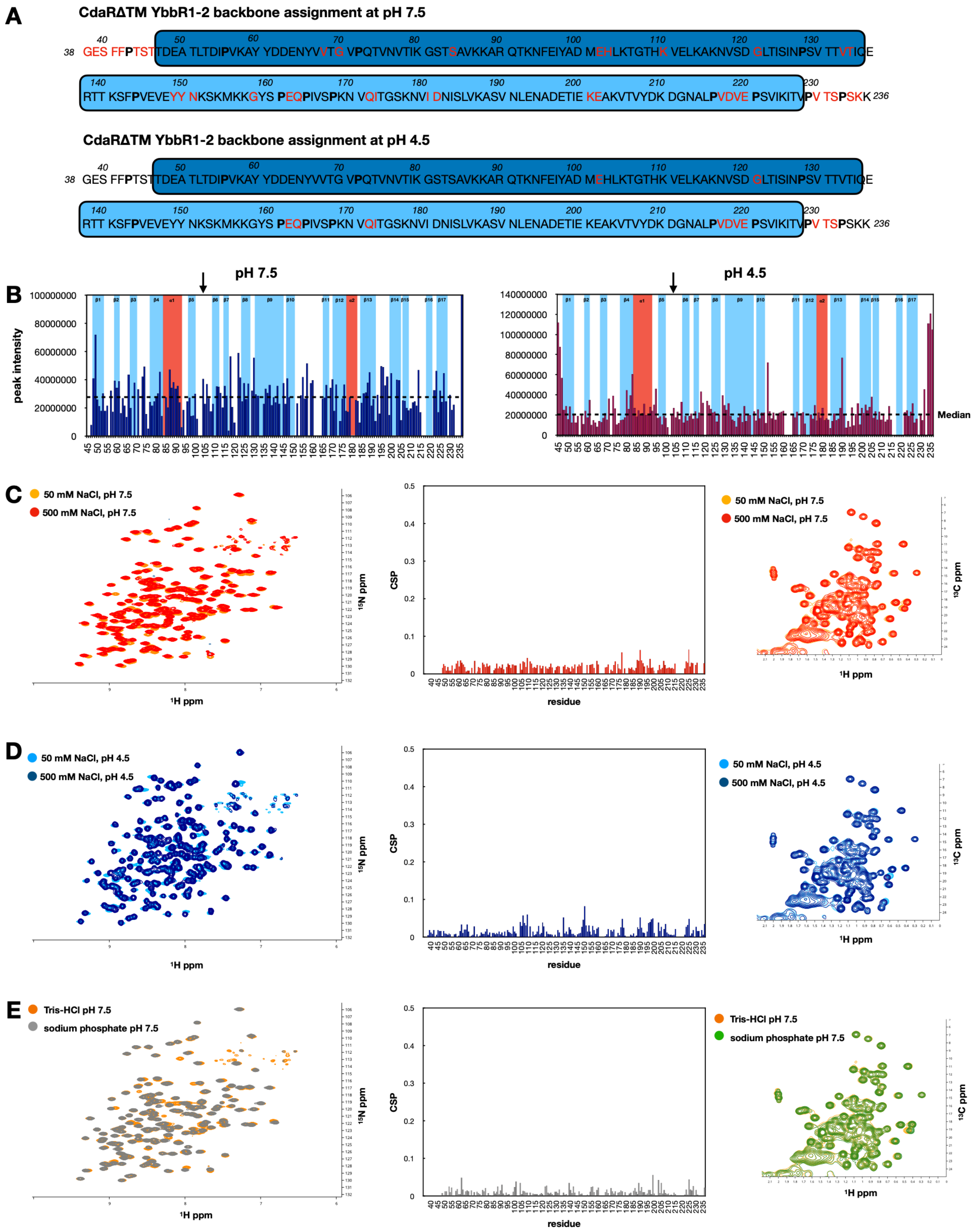


**Fig. S6.**

**A)** Backbone assignment of *Bacillus subtilis* CdaRΔTM YbbR1-2 at pH 7.5 (top) and pH 4.5 (bottom). Residues that could not be assigned are shown in red, prolines in bold and YbbR domain

1 in dark blue and 2 in light blue. **B)** NMR peak intensities at pH 7.5 and pH 4.5, showing a lower peak intensity around the unassigned residue E102 (black arrow). β-sheets are marked in light blue and $\alpha$-helices in red. **C)** The effect of salt on $^{1}H$-$^{15}N$ (left) and $^{1}H$-$^{13}C$ (right) chemical shifts in CdaRΔTM YbbR1-2. in pH 7.5 The $^{1}H$-$^{15}N$ chemical shift changes are plotted against the sequence in the middle graph. **D)** Same experiment as C but at pH 4.5. **E)** The effect of the buffer used on chemical shift changes.

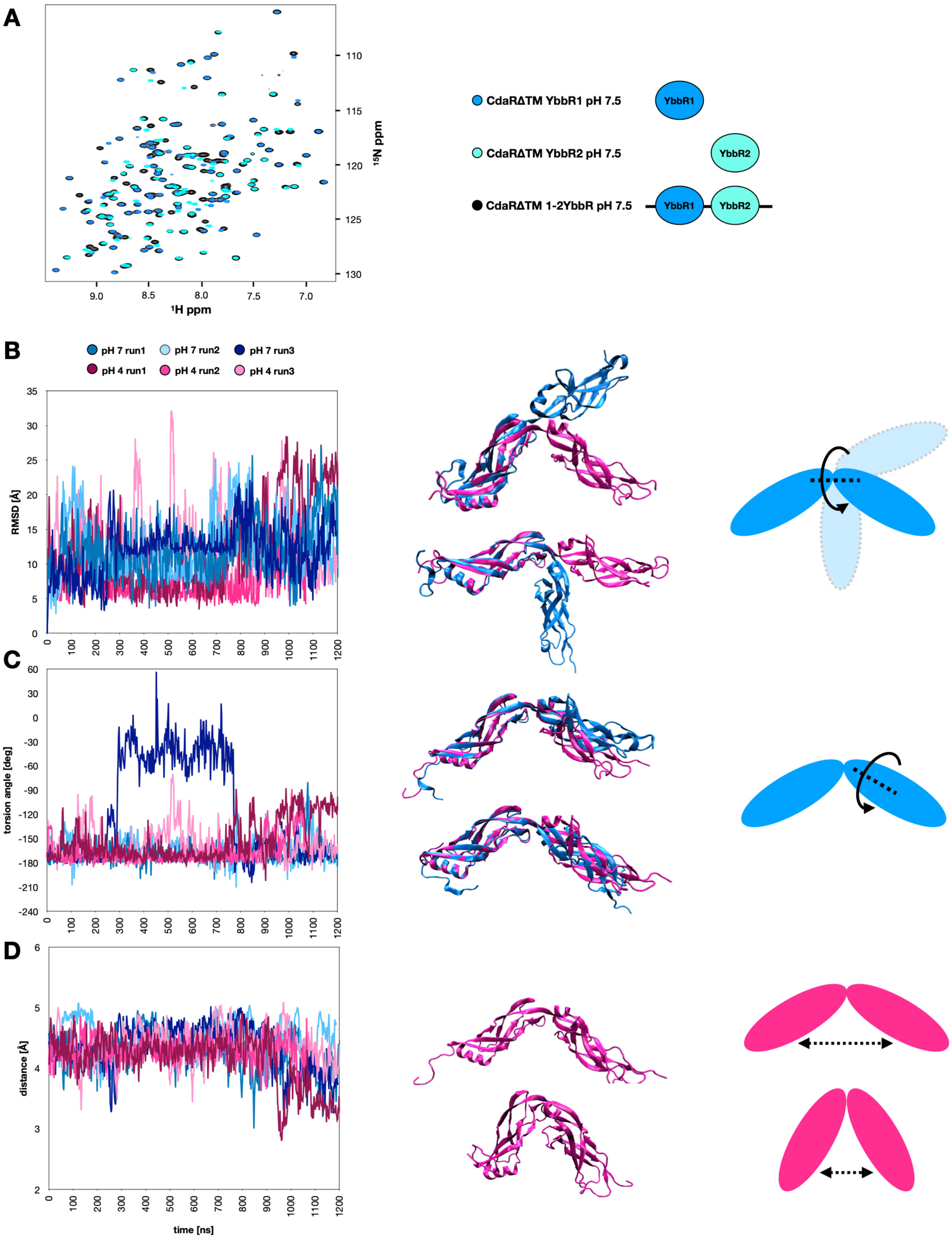


**Fig. S7.**

**A)** $^{1}$H-$^{15}$N HSQC spectra of CdaRΔTM YbbR1-2 (black), CdaRΔTM YbbR1 (light blue) and CdaRΔTM YbbR2 (cyan) at pH 7.5. The peak position in both the individual and combined constructs overlay perfectly, with exception of the N-terminal regions of the constructs that carry the affinity tag. **B)** RMSD at pH 7 and pH 4 during the trajectory compared to its original position. On the right examples of the different positions YbbR domain 1 can adopt with respect to YbbR domain 2, showing high flexibility of the two domains with respect to each other. **C)** Left: Torsion angle between YbbR domain 1 and 2 during the trajectory. In the middle, visualisation of the twisting movement performed by YbbR domain 2 at pH 7 run1, seen as a large change in torsion angle between 300 and 800 ns. **D)** The change in distance between the 2 domains over the trajectory at the 2 pHs. The closest distance can be observed at pH 4 run1 at the end of the trajectory, resulting in a “closing” of the two domains (right).

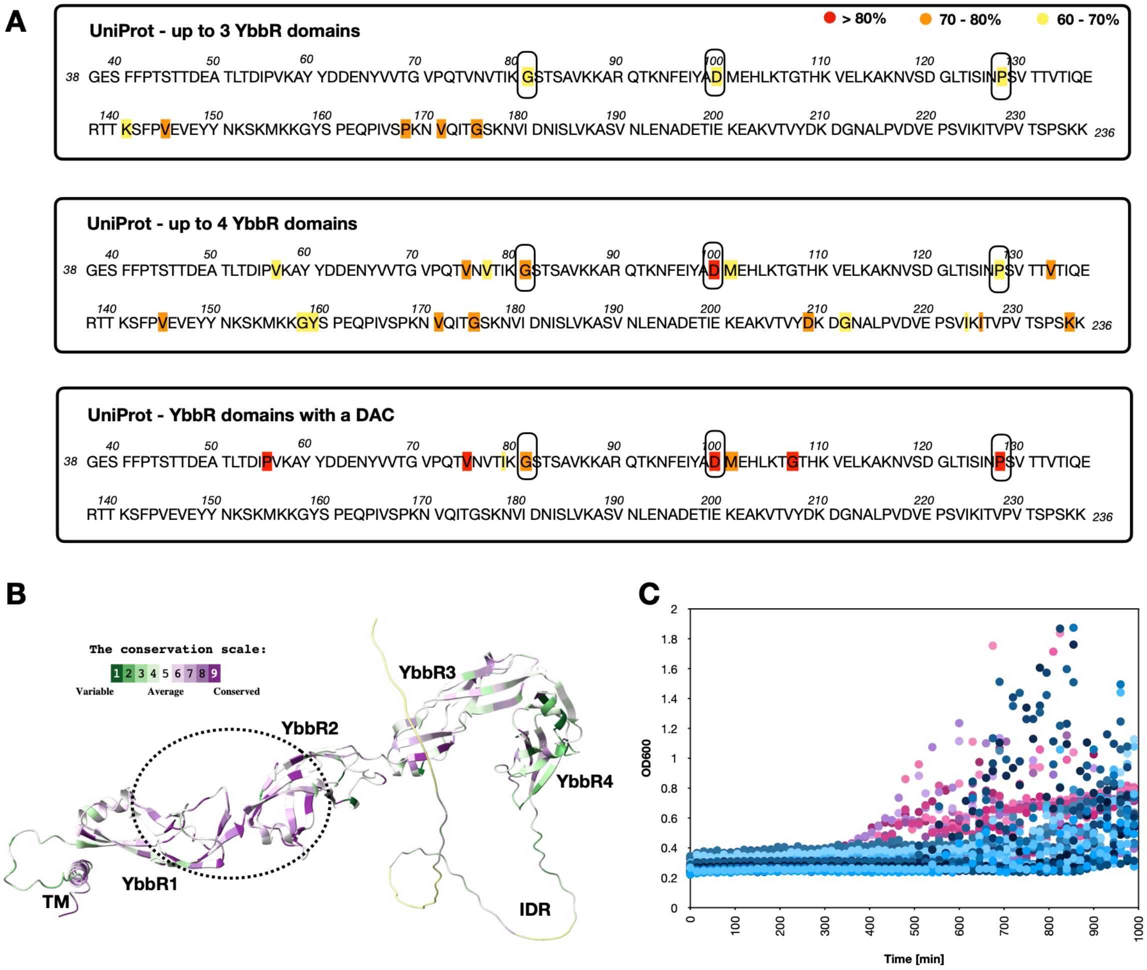


### Fig. S8.

**A)** Multiple sequence alignment using ClustalΩ and different search options for YbbR domains in the Uniprot database. Residues that are conserved in all sequence alignments are circled with a black line. **B)** Evolutionary conservation of residues in the *B. subtilis* CdaR protein using the software tool ConSurf. The highlighted hinge region between domain 1 and 2 is highly conserved. **C)** Growth curves of WT *B. subtilis* (blue) and the D100A mutant (pink) in $LB_{BS}$ pH 4.5. Shown are 7 independent growth experiments including three clones each carrying the CdaR D100A mutation. Of note the high dispersion of the data points due to the formation of filaments at low pH (see Fig. S1A).

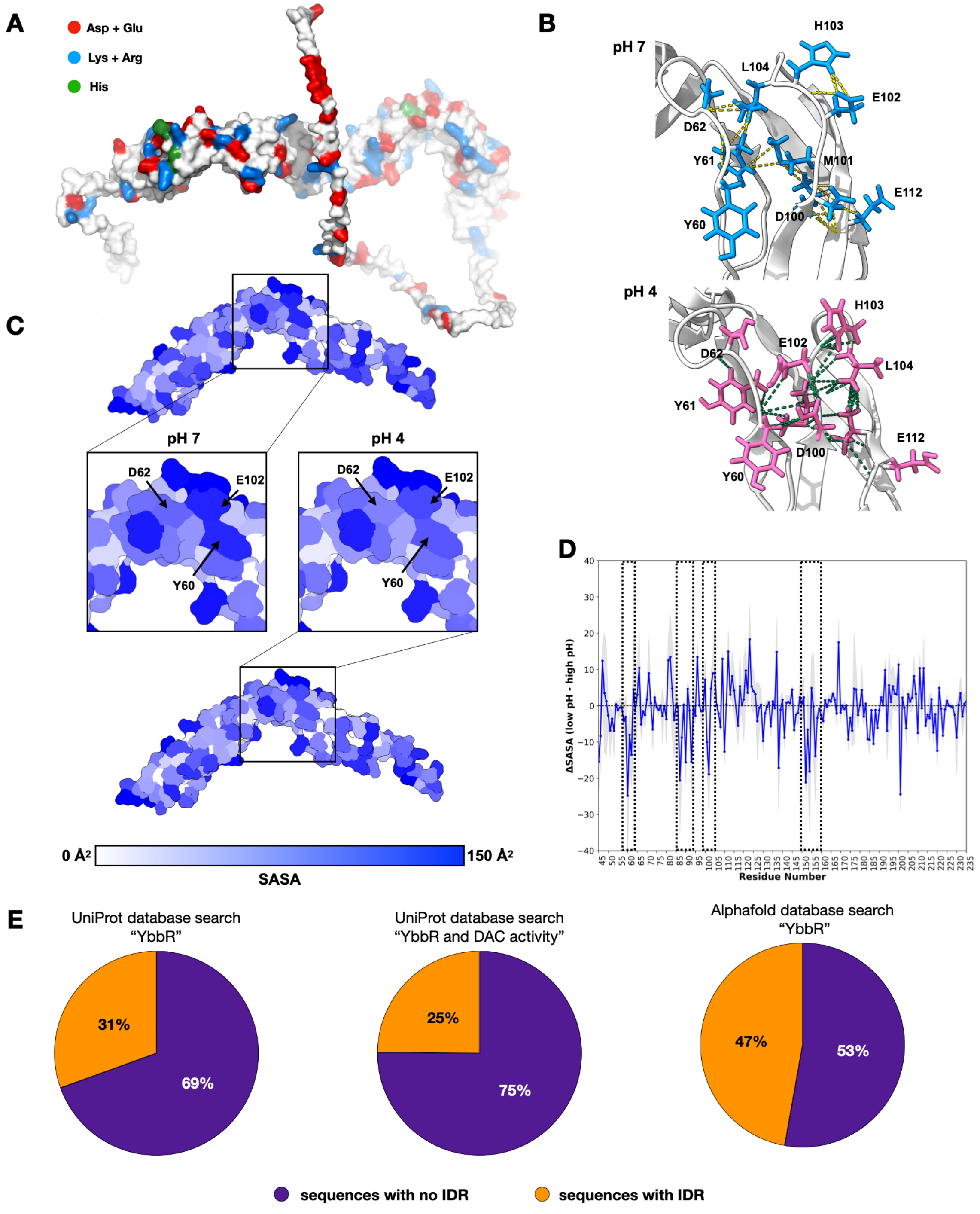


**Fig. S9.**

**A)** Charge distribution in the CdaR protein. As can be seen charges are distributed evenly over the protein with no clustering in the hinge region between domain 1 and 2. **B)** Change of the interaction networks of the residues in the hinge region at pH 7 (top, blue) and pH 4 (bottom, pink). Interactions are shown using the ChimeraX contacts tool with VDW overlap ≥ -0.4 Å. **C)** Comparison of the solvent-accessible surface area (SASA) during MD simulations at pH 7 (top, middle left) and pH 4 (bottom, middle right). Solvent exposure (solvent-accessible surface area - SASA) is depicted in a gradient from white to blue, with the highest solvent exposure depicted in dark blue. **D)** Residuewise difference in SASA between low-pH and high-pH simulations. Average per-residue SASA values were calculated from molecular dynamics trajectories and compared between conditions as ΔSASA. Positive values indicate increased solvent exposure at low pH, whereas negative values indicate reduced exposure. The solid blue line represents the mean ΔSASA across replicate simulations, and shaded grey regions indicate the standard deviation. regions that show the highest solvent protection at low pH are highlighted by a dashed line, these include: the hinge region around D100 (E102, M101, Y60, Y61 and D62) a helix in YbbR domain 1 around residue 90 and the helix formed at low pH around residue 155. **E)** Prediction of IDRs using the software ALBATROSS on sequences deposited in the UniProt or AlphaFold database containing YbbR domains or YbbR domains and DAC catalytic activity. In all searches more than half of the sequences carry no IDR.

**Table S1. X-ray crystallography statistics of PDB 32QW and 32RB.**

| | pH 7.5 | pH 5.0 |
|---|---|---|
| **Resolution range** | 36.38 - 1.16 (1.19 - 1.16) | 26.25 - 1.12 (1.14 - 1.12) |
| **Space group** | C 1 2 1 | C 1 2 1 |
| **Unit cell** | 72.36 45.35 148.78 90 101.99 90 | 71.91 45.93 148.63 90 101.1 90 |
| **Unique reflections** | 144275 (2670) | 136965 (368) |
| **Completeness (%)** | 88.41 (22.99) | 74.26 (2.80) |
| **Wilson B-factor** | 17.72 | 17.45 |
| **Reflections used in refinement** | 144275 (2670) | 136965 (368) |
| **Reflections used for R-free** | 1903 (39) | 1928 (7) |
| **R-work** | 0.17 (0.43) | 0.19 (0.43) |
| **R-free** | 0.21 (0.50) | 0.21 (0.39) |
| **Number of non-hydrogen atoms** | 3907 | 3811 |
| macromolecules | 3227 | 3143 |
| solvent | 680 | 668 |
| **Protein residues** | 403 | 402 |
| **RMS(bonds)** | 0.006 | 0.006 |

| | | |
|---|---|---|
| **RMS(angles)** | 0.86 | 0.83 |
| **Ramachandran favored (%)** | 98.75 | 99.25 |
| **Ramachandran allowed (%)** | 1.25 | 0.75 |
| **Ramachandran outliers (%)** | 0.00 | 0.00 |
| **Rotamer outliers (%)** | 0.79 | 0.00 |
| **Clashscore** | 0.93 | 1.11 |
| **Average B-factor** | 28.71 | 23.07 |
| macromolecules | 26.07 | 21.73 |
| solvent | 41.25 | 29.39 |

**Movie S1 (separate file).** Formation of an alpha-helix at low pH during the MD simulation.

**Movie S2 (separate file).** Unfolding of the hinge region at high pH during the MD simulation.

**Movie S3 (separate file).** Motion of the hinge region at low pH during the MD simulation.

**Movie S4 (separate file).** 180º flip of Ybbr domain 2 at high pH during the MD simulation.

**Movie S5 (separate file).** “Closing” of Ybbr domain 1 and 2 at low pH during the MD simulation.
